# Canonical AREs are tumor suppressive regulatory elements in the prostate

**DOI:** 10.1101/2024.02.23.581466

**Authors:** Michael A. Augello, Xuanrong Chen, Deli Liu, Kevin Lin, Alex Hakansson, Martin Sjöström, Francesca Khani, Lesa D. Deonarine, Yang Liu, Jaida Travascio-Green, Jiansheng Wu, Massimo Loda, Felix Y. Feng, Brian D. Robinson, Elai Davicioni, Andrea Sboner, Christopher E. Barbieri

**Affiliations:** Department of Urology, Weill Cornell Medicine, New York, NY 10065, USA; Sandra and Edward Meyer Cancer Center, Weill Cornell Medicine, New York, NY 10065, USA; The HRH Prince Alwaleed Bin Talal Bin Abdulaziz Alsaud Institute for Computational Biomedicine, Weill Cornell Medicine, New York, NY 10065, USA; Veracyte, Inc., South San Francisco, CA 94080, USA; Department of Radiation Oncology, University of California, San Francisco, CA 94115; Department of Pathology and Laboratory Medicine, Weill Cornell Medicine, New York, NY 10065, USA; Departments of Urology and Medicine University of California, San Francisco, CA 94115; Caryl and Israel Englander Institute for Precision Medicine, Weill Cornell Medicine, New York, NY 10065, USA

## Abstract

The androgen receptor (AR) is the central determinant of prostate tissue identity and differentiation, controlling normal, growth-suppressive prostate-specific gene expression^1^. It is also a key driver of prostate tumorigenesis, becoming “hijacked” to drive oncogenic transcription^2-5^. However, the regulatory elements determining the execution of the growth suppressive AR transcriptional program, and whether this can be reactivated in prostate cancer (PCa) cells remains unclear. Canonical androgen response element (ARE) motifs are the classic DNA binding element for AR^6^. Here, we used a genome-wide strategy to modulate regulatory elements containing AREs to define distinct AR transcriptional programs. We find that activation of these AREs is specifically associated with differentiation and growth suppressive transcription, and this can be reactivated to cause death in AR^+^ PCa cells. In contrast, repression of AREs is well tolerated by PCa cells, but deleterious to normal prostate cells. Finally, gene expression signatures driven by ARE activity are associated with improved prognosis and luminal phenotypes in human PCa patients. This study demonstrates that canonical AREs are responsible for a normal, growth-suppressive, lineage-specific transcriptional program, that this can be reengaged in PCa cells for potential therapeutic benefit, and genes controlled by this mechanism are clinically relevant in human PCa patients.

## Main

Many cancers show dependence on tissue and lineage-specific transcription factors that are critical for normal tissue as well^7^. The androgen receptor (AR) is the central determinant of prostate tissue identity, lineage, and differentiation, controlling normal, growth-suppressive prostate-specific gene expression^1^. However, it is also a key driver of prostate tumorigenesis, becoming “hijacked” through epigenetic reprogramming to drive oncogenic transcription^3-5,8^. Importantly, it remains the key therapeutic target for PCa, even in advanced, treatment-resistant disease^9^, where genomic alterations such as *AR* gene and regulatory element amplification, overexpression, mutations, and splice variants of *AR* drive continued reliance on androgen signaling^10^.

Most focus in the field of AR reprogramming has been on gain of oncogenic functions by AR, associated with a shift in cistromic localization and control of new target genes associated with oncogenic effects such as proliferation and invasion^3,11^. However, reprogramming in both directions may be important – loss of tumor suppressive functions may be critical. Importantly, evidence suggests oncogenic and growth-suppressive transcriptional programs controlled by AR are associated with distinct epigenomic regulation. Classically, nearly all AR-regulated gene expression is mediated through direct AR binding to palindromic DNA sequences known as Androgen Response Elements (AREs)^1^. In PCa, the AR cistrome is distinct and highly associated with motifs for FOXA1 and HOXB13, and these can drive the AR cistrome to reflect PCa^3,12-14^, while other genomic alterations impair the normal growth suppressive cistrome and transcriptome of AR^4^, revealing the plasticity and context-dependency of AR programs.

However, several major issues remain largely unexplained: What is the importance of canonical AREs in differentiating oncogenic and growth-suppressive AR-driven transcriptional programs? Can these programs be separated and independently modulated? Do canonical AREs remain essential for PCa cells? We, therefore, sought to better characterize the epigenomic regulation of the growth suppressive AR program, its control by AREs, and its deregulation in human PCa.

### AREs are enriched in the AR cistrome of normal prostate tissue and depleted in prostate cancer

In cohorts of human normal prostate tissue and PCa samples^3,8,15^, AREs were less common in the AR cistrome of PCa compared to normal prostate tissue (**Fig. 1a**). Further interrogation in a dataset of matched tumor and normal tissue from the same patient demonstrated that each patient sample showed depletion of AREs in tumor compared to matched normal^15^ (**Fig. 1b**). Enhancer activity (H3K27ac ChIP-seq) confirms that AR binding patterns reflect active regulatory control at tumor- and normal-specific AR peaks (**Fig. 1c**). Finally, ectopic expression of AR reprograming factors associated with PCa (FOXA1 and HOXB13) in non-transformed prostate cells^3^ resulted in depletion of AREs from the AR cistrome (**Fig. 1d**). Therefore, ARE motifs were associated with normal prostate tissue and de-enriched in PCa, suggesting a tumor-suppressive regulatory element. We therefore hypothesized that activation of ARE-containing regulatory elements would have tumor suppressive effects in prostate cells.

**Figure 1.**
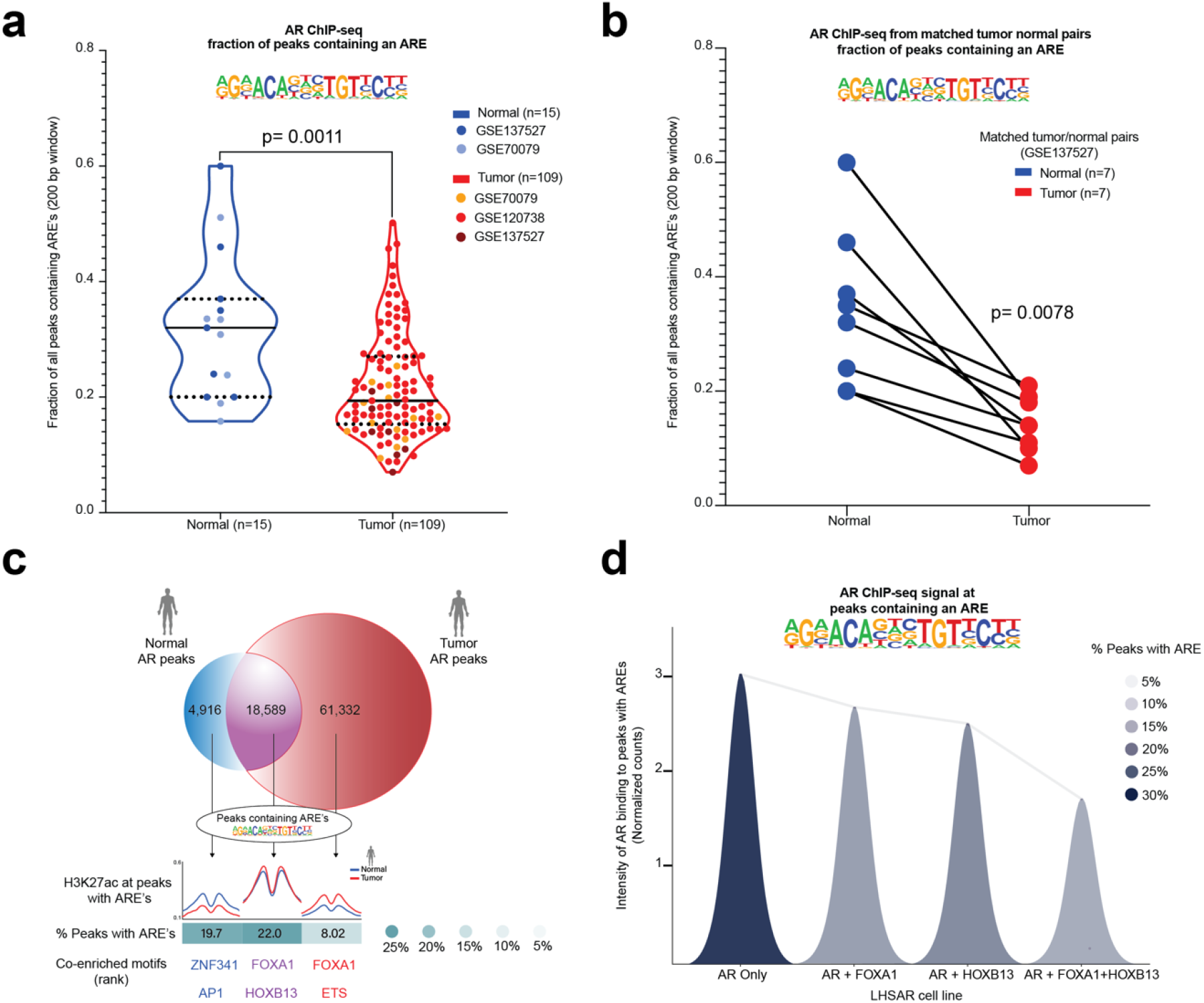
Canonical AR Response Element (ARE) motifs are depleted in human prostate cancer. **a**, Fraction of peaks containing an ARE from AR ChIP-seq data in normal human prostate tissue (blue) and PCa tissue (red). *P* =□0.0011 by Mann-Whitney two-sided. **b**, Fraction of peaks with ARE from AR ChIP-seq data in matched normal human prostate tissue (blue) and PCa (red) from the same patients. *P*□=□0.0078 by Wilcoxon two-sided test. **c**, Overlap of collated AR peaks from normal and tumor prostate samples, with associated enhancer activity (H3K27ac) at normal specific (blue), tumor specific (red), and common (purple) AR peaks, along with percent containing AREs, and other associated motifs. **d**, Intensity of AR binding in peaks with an ARE in LHSAR cells expressing AR alone or associated oncogenic factors (FOXA1 and HOXB13). AR binding to peaks with AREs decreases with addition of FOXA1 and HOXB13.

### Development and validation of a novel strategy to modulate ARE-associated regulatory elements

To test this, we developed inducible constructs to modulate chromatin activity around AREs. ***MACCs*** (**M**odifiers of **A**RE **C**ontaining **C**hromatin – **Fig. 2a**) are designed to localize to AREs via the DNA binding domain of AR, but lack the N-terminal region largely responsible for recruitment of co-factors^16^ (Methods and Extended Data Fig. 1). MACCs are tamoxifen-inducible, epitope-tagged (3X FLAG), and contain repressive or activating chromatin modifying domains (H3K9 methyltransferase KRAB, or transcriptional activator VP64). We examined localization and activity of three different MACCs, named regarding expected transcriptional effects: 1) Neutral **(N**, 3X FLAG alone, no chromatin modifying domain), 2) Repressive (**R**, KRAB), 3) Activating (**A**, VP64). In stable LNCaP lines, after confirming expression and tight inducible nuclear localization with tamoxifen (**Fig. 2a, b**), we examined genome-wide distribution with ChIP-seq. Consistent with appropriate localization, consensus MACC peaks (Extended Data Fig. 2a,b) localized primarily to introns and intergenic regions (Extended Data Fig. 2c), and ARE was the most significantly enriched motif, followed by lower affinity variants of the ARE such as AR-halfsites^17,18^ (**Fig. 2c**). Modulation of H3K27ac was consistent with expected activity – at ARE-containing peaks, H3K27ac was high with activating MACC, and lowest with repressive (**Fig. 2c**, Extended Data Fig. 2d). The well-characterized AR-target gene *FKBP5* provides a clear example, with three enhancer elements (E1, E2, and E3) bound by AR^19^. E1 and E2 contain AREs, are bound by MACC constructs, and show expected modulation of H3K27ac, while E3 lacks a canonical ARE motif, and shows no MACC effects on H3K27ac (**Fig. 2d**). Further, MACC activity was restricted to AR bound chromatin enriched for AREs (Extended Data Fig. 2e), as other AR binding sites lacking an ARE showed no global difference in H3K27ac signal (Extended Data Fig. 2e). Finally, interrogation of H3K27ac ChIP-seq data from human patient samples^20^ showed that enhancer activity at MACC consensus peaks distinguishes tumor from normal tissue (**Fig. 2e)**, highlighting the relevance of the binding sites of these engineered constructs in human prostate tissue. These MACC constructs represent novel tools to modulate ARE-containing regulatory elements, reveal and manipulate distinct subsets of AR-responsive enhancers, and interrogate cancer and normal-specific signaling pathways, with clear applicability to human specimens.

**Figure 2.**
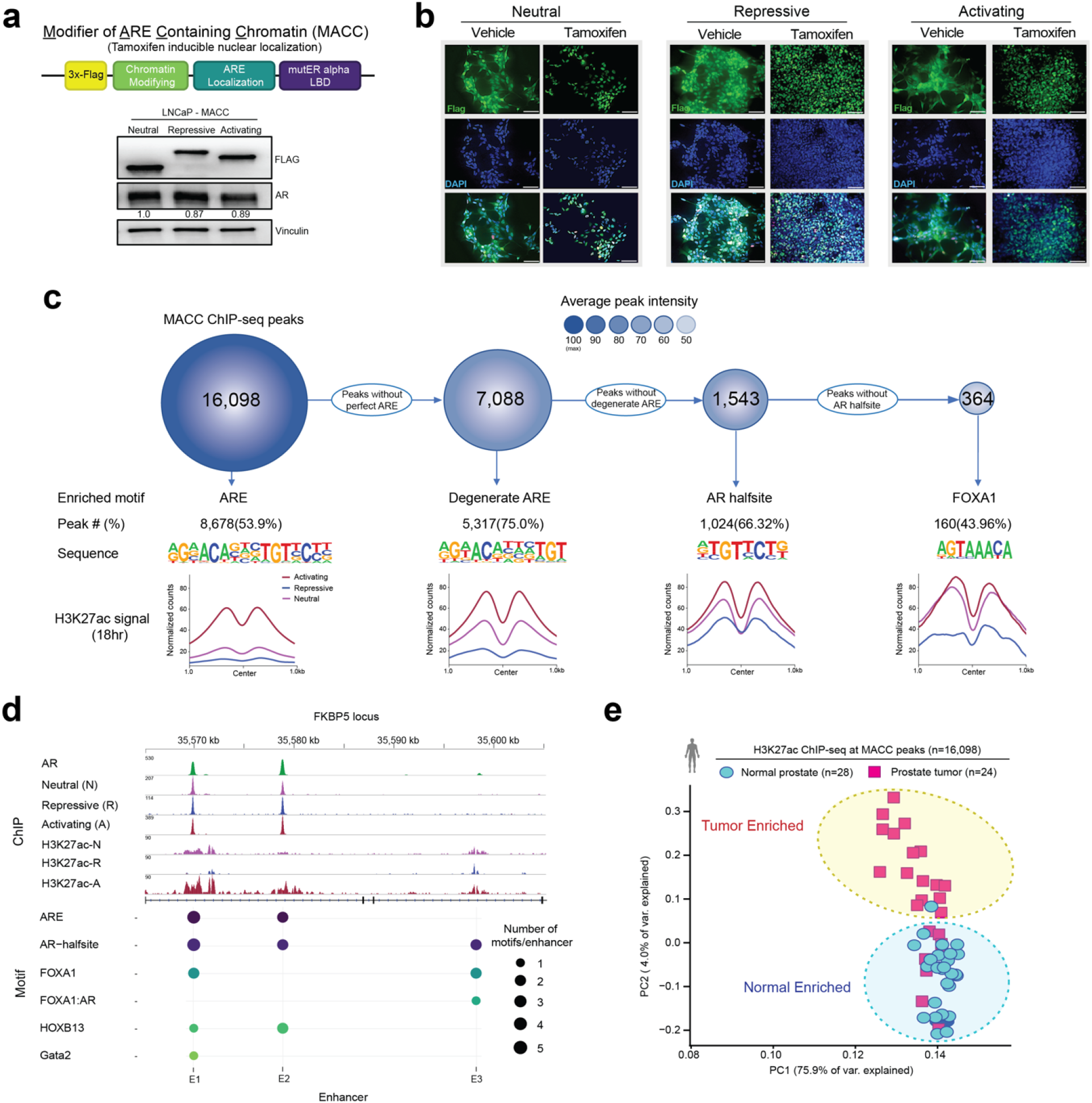
MACCs represent an inducible system to directly modulate ARE-containing regulatory elements. **a**, Schematic of MACC constructs, and expression of neutral (3X FLAG only), repressive (KRAB) and activating (VP64) constructs. **b**, Nuclear localization of constructs upon tamoxifen induction. Scale bar = 20□μm. **c**, ChIP-seq of MACC constructs in LNCaP cells, showing maximal binding and effect on enhancer activity at ARE motifs, with decreasing affinity and effect on H3K27ac activity for associated motifs. **d**, Example of MACC localization and modulation of regulatory activity at the FKBP5 locus, with ARE-containing enhancers 1 and 2 (E1 and E2) affected by MACCs, but E3 (without an ARE) is insensitive. **e**, H3K27ac signal at MACC peaks distinguishes tumor from normal human prostate tissue.

### Activation of ARE-enriched enhancers is growth suppressive in prostate cancer cells in vitro and in vivo and dispensable for tumorigenic phenotypes

To determine the biological impact of directly modulating these ARE-containing regulatory elements, we examined the effect of induction of MACC constructs in androgen-dependent PCa cell lines. Dogma states that PCa cells respond to AR activation with concomitant increased expression of AR target genes and increased proliferation^21^, and that these are tightly coupled. To clarify the effects on the transcriptional programs controlled specifically by AREs in PCa cells, we performed gene expression profiling with RNA-seq in LNCaP with tamoxifen-inducible activation of MACC constructs (**Fig. 3a,b**), principle component analysis showed that without induction (vehicle) alone, all LNCaP lines had similar transcriptomes (**Fig. 3a**). In contrast, induction with tamoxifen led to dramatic changes in transcriptomes with activating and repressive MACCs (**Fig. 3a**). Pathway analysis showed that activation of AREs resulted in upregulation of classic AR target genes (GSEA AR Hallmarks^22^), with simultaneous downregulation of gene sets associated with proliferation (**Fig. 3b**). These results were further confirmed by analyzing clinically relevant transcriptional signatures of AR (AR score)^23^ and the cell cycle (RB loss signature)^24^ (**Fig. 3c**). Activation of AREs was associated with a higher AR score but was de-enriched cell cycle activity. Conversely, ARE repression resulted in an inverted signature, with a lower AR score and higher cell cycle de-regulation signature (**Fig. 3b,c)**. Collectively, these data suggest that there is a decoupling of AR activity from cell cycle effects by modulating enhancers enriched for AREs.

**Figure 3.**
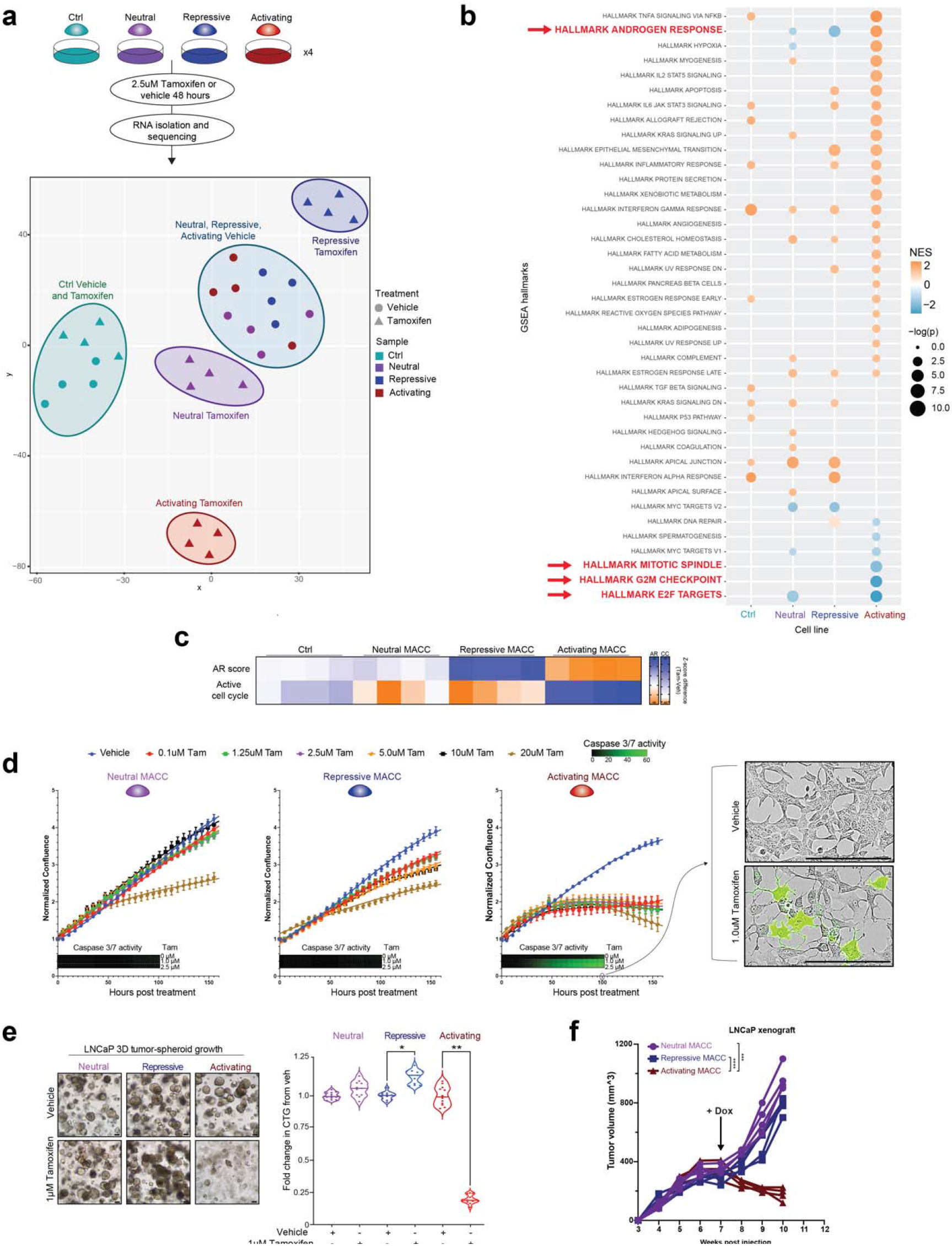
Modulation of AREs results in uncoupling of canonical AR target genes and proliferation in prostate cancer cells, with ARE activation growth suppressive, while ARE repression is tolerated. **a**, Experimental plan for transcriptional profiling, and PCA of gene expression profiled from vehicle or tamoxifen treated LNCaP MACC lines. **b**, Pathway analysis using GSEA Hallmarks of induced vs. vehicle LNCaP MACC lines. **c**, Opposing effects on AR target genes (AR score) and cell cycle genes (RB loss signature) with activation or repression of AREs. **d**, Growth of LNCaP MACC lines in 2D culture with vehicle or increasing doses of tamoxifen. Neutral and repressive constructs have minimal effect; Activating MACC is growth suppressive and induces apoptosis as measured by Caspase 3/7 activity. Scale bar = 20□μm. **e**, Brightfield images and growth of LNCaP MACC lines as 3D spheroids, +/- tamoxifen. Scale bar = 20□μm. Two-tailed Student’s t-test, ^*^*p*□<□0.05, ^**^*p*□<□0.01. **f**, LNCaP xenografts with doxycycline-inducible MACC constructs in nude mice. Dox chow started at week 7. Two-way ANOVA, ^***^*p*□<□0.001, ^****^*p*□<□0.0001.

In stable LNCaP lines with inducible expression of **N**eutral, **R**epressive, and **A**ctivating MACCs, activation of AREs resulted in severe growth suppression and cell death in both basal growth conditions and androgen-simulated growth after androgen starvation (**Fig. 3d** and Extended Data Fig. **3**). Repression of AREs had minimal effect in 2D culture (**Fig. 3d** and Extended Data Fig. **3**), but stimulated increased growth of LNCaP cells as 3D spheroids (**Fig. 3e**). Similar effects were observed in another androgen-dependent prostate cell line (LAPC4, Extended Data Fig. 4a-d), but non-prostate cells (293T), and androgen-indifferent PCa cells (PC3, DU145) showed no effect of MACC induction (Extended Data Fig. 4e-g), consistent with lineage specificity. To confirm that effects were not artifact related to hormonal treatment with tamoxifen, we engineered an additional doxycycline-inducible system with the same biological effects (Extended Data Fig. 4a). Finally, activation of AREs in vivo also suppressed the growth of LNCaP xenografts in nude mice, while ARE repression was dispensable for tumor growth (**Fig. 3e** and Supplementary Fiig.1). Together, these data show that direct activation of regulatory elements containing AREs results in growth suppression and cell death of PCa cells.

**Figure 4.**
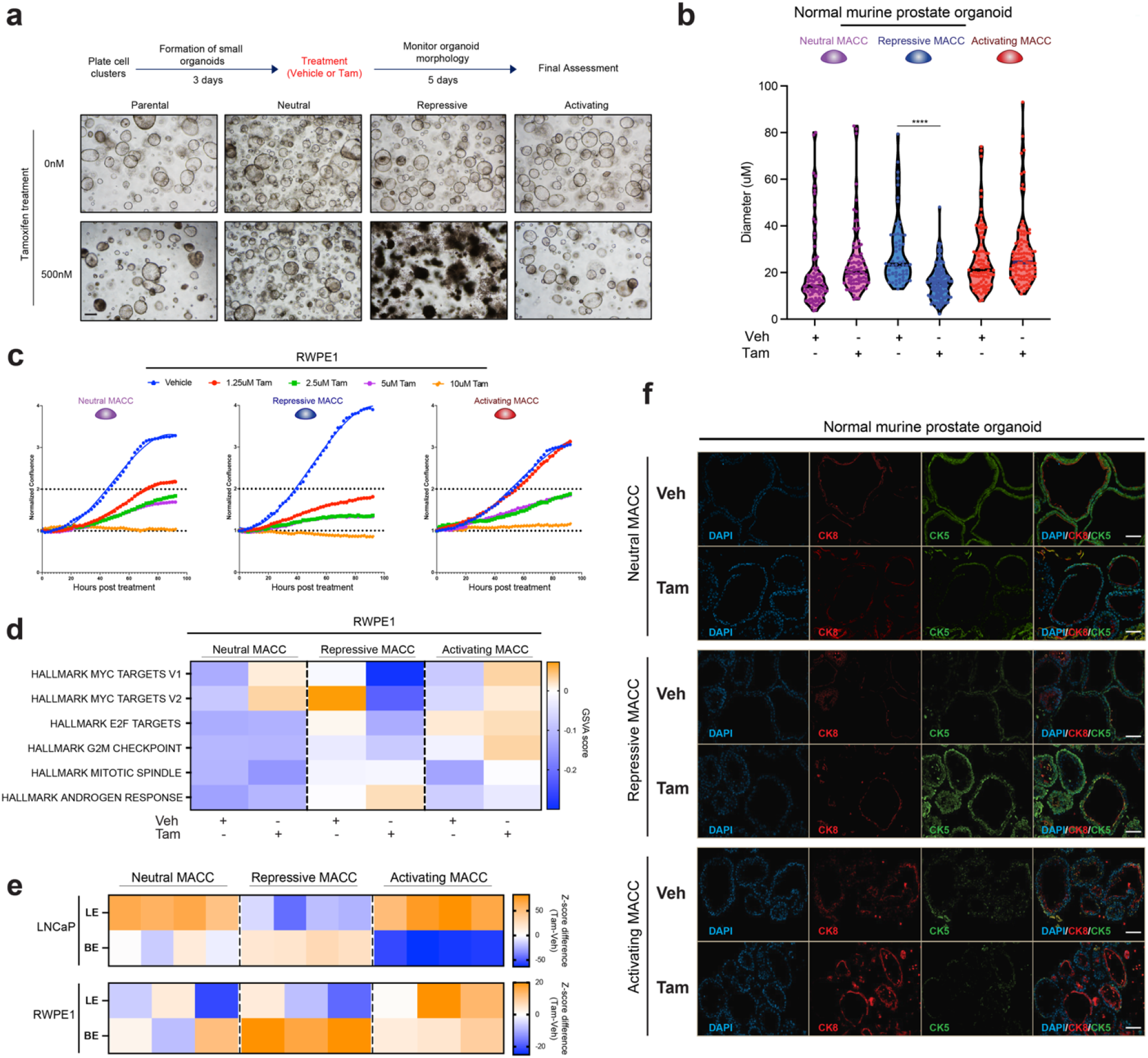
In benign prostate cells, ARE activation is well tolerated, but ARE repression results in altered growth and differentiation. **a-b**, Genetically normal mouse prostate organoids (**a**, #1 and **b**, #2) with inducible MACC constructs, showing loss of luminal morphology with repression of AREs. Scale bar = 40□μm. Two-tailed Student’s t-test, ^****^*p*□<□0.0001. **c**, Growth of benign RWPE1 prostate MACC lines with vehicle or increasing doses of tamoxifen. **d**, Pathway analysis using gene expression profiling of induced vs. vehicle RWPE MACC lines. **e**, Heatmap of luminal epithelial (LE) and basal epithelial (BE) markers’ gene signature levels in LNCaP and RWPE1 MACC lines. **f**, Representative immunofluorescence images of the expression of luminal (cytokeratin 8) and basal (cytokeratin 5) markers in normal mouse prostate organoids (#2). Organoids were treated with either vehicle or 2uM Tamoxifen for 48h. Scale bar = 20□μm.

### Repression of ARE-associated chromatin disrupts differentiation in normal prostate epithelial cells

Our data in PCa cell lines showed that activation of AREs results in growth suppressive phenotypes, while repression of AREs has minimal effects. We next sought to determine the effects of manipulating these regulatory elements in normal prostate epithelial cells by deploying inducible MACC constructs in genetically normal mouse prostate organoids. Neutral and activating MACCs had no distinguishable effect on organoid phenotypes, while the repressive MACC resulted in severe disruption of growth and luminal features in multiple independent mouse organoid lines (**Fig. 4a,b** and Extended Data Fig. 5a-d). In direct contrast to PCa cell lines, repression of AREs in human benign prostate epithelial cells (RWPE1) resulted in growth suppression, while activation of AREs had fewer effects (**Fig. 4c** and Extended Data Fig. 5e,f). Gene expression profiling in these non-transformed prostate cells was consistent with these effects, with downregulation of gene sets associated with proliferation (e.g. Myc and E2F targets, G2M checkpoint) and ARE repression (**Fig. 4d**, Extended Data Fig.6 and Supplementary Fiig.2). In addition, modulation of AREs revealed regulation of prostate epithelial differentiation phenotypes. In both LNCaP and RWPE1 cells, ARE repression led to downregulation of genes associated with luminal epithelia (LE), and upregulation of basal epithelial (BE) markers, while ARE activation maintained luminal gene expression (**Fig. 4e**). Finally, in normal mouse prostate organoids, immunofluorescence showed altered expression of luminal (cytokeratin 8) and basal (cytokeratin 5) markers with ARE modulation (**Fig. 4f**). These data show that in normal prostate epithelial cells, elevation of transcriptional programs controlled by AREs has minimal effects, while disruption of these regulatory elements results in growth suppression and loss of luminal epithelial phenotypes.

**Figure 5.**
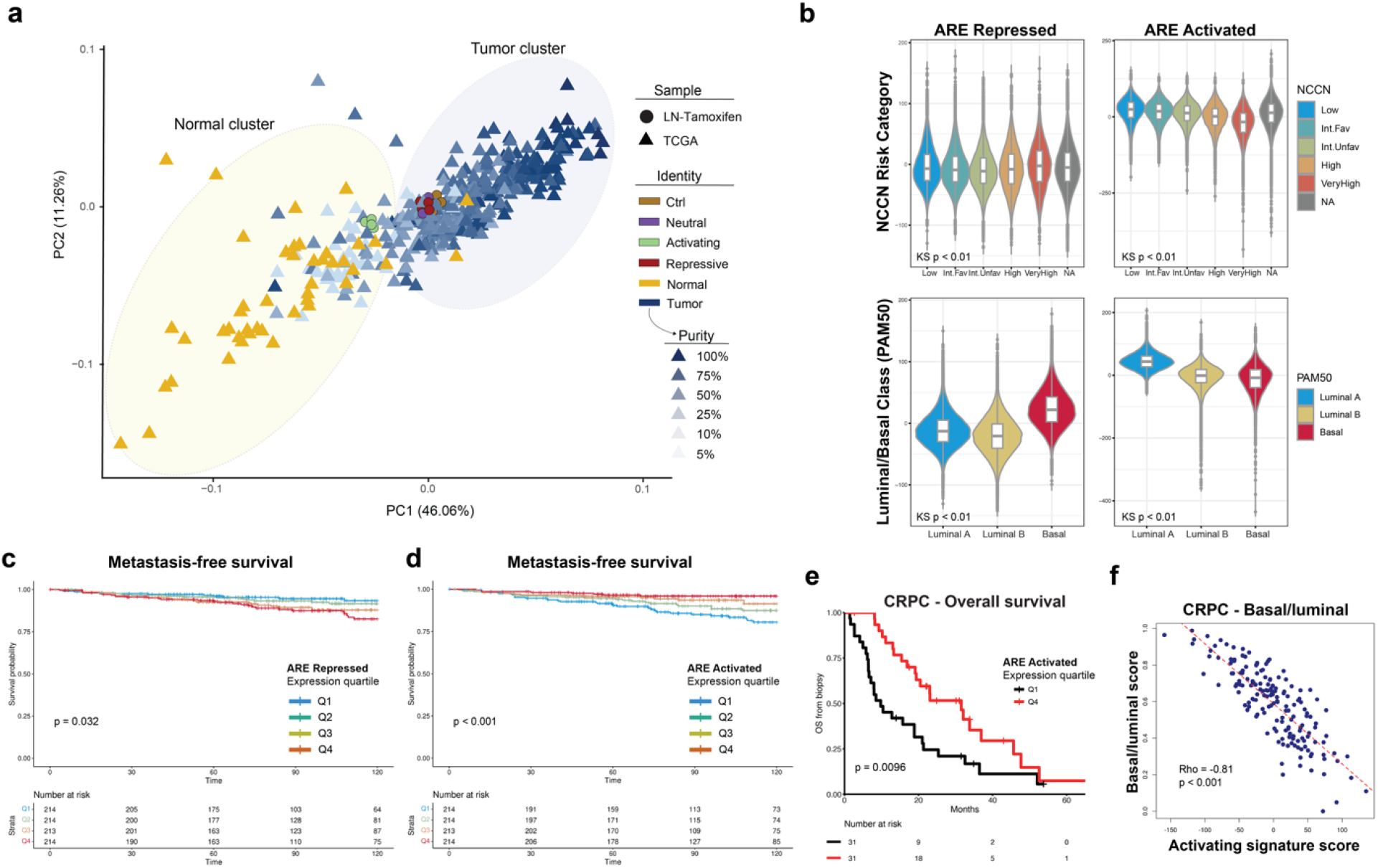
Distinct transcriptional programs revealed by modulating AREs are clinically relevant in human prostate cancer. **a**, Unsupervised clustering of the transcriptomes of human prostate cancer and normal samples (TCGA), along with tamoxifen induced LNCaP MACC lines. **b**, Association of ARE and activated signatures from LNCaP MACC lines with National Comprehensive Cancer Network (NCCN) risk category (top) and PAM50 luminal/basal classification (bottom) in clinically localized prostate cancer, n=169,123. KS = Kruskal-Wallis. Int.Fav = Favorable Intermediate risk. Int.Unfav = Unfavorable Intermediate risk, also Supplementary Table 1 and 3. **c**, Kaplan Meier analysis of metastasis-free survival in 855 men after radical prostatectomy, stratified into quartiles by ARE repressed signature expression level. Time = months. **d**, Kaplan Meier analysis of metastasis-free survival in 855 men after radical prostatectomy, stratified into quartiles by ARE activated signature expression level. Time = months. **e**, Kaplan Meier analysis of overall survival in 123 men with metastatic castrate resistant prostate cancer, stratified into quartiles by ARE activated signature expression level in biopsy specimens. Hazard ratio (HR) = 0.47 [0.26−0.83]. **f**, Correlation analysis of basal/luminal phenotype score (higher = more basal) by gene expression (Y-axis) with ARE activated signature expression level (X-axis). Rho=-0.81, Spearman’s correlation.

### Distinct transcriptional programs revealed by modulating AREs are clinically relevant in human prostate cancer

We next asked whether the distinct AR target genes revealed by our strategy to directly modulate AREs were relevant in human prostate samples. We examined the relationship of transcriptional programs controlled by AREs to human PCa and normal prostate tissue samples. Unsupervised clustering of the transcriptomes of LNCaP cells with inducible expression of MACCs with human PCa and normal prostate samples^25^ revealed that all samples clustered with tumor samples except those with activation of AREs, whose transcriptional profile shifted toward normal prostate tissue (**Fig. 5a** and Extended Data Fig. 7a). Similar effects were observed in an independent cohort with matched cancer and normal tissue^26^ (Extended data Fig.7b,c). We use the transcriptional responses of LNCaPs to generate gene expression signatures specific to each MACC construct, i.e., ARE activation or repression (Extended Data Fig.8). Application of these signatures to single-cell data from human PCa specimens^27^ showed enrichment of the signature of ARE activation primarily in *normal* luminal epithelial cells, while the signature of ARE repression was enriched in *tumor* luminal epithelium (Extended Data Fig. 9). We next examined the impact of these signatures on the prognosis in patients with PCa. In a cohort of over 169,000 patients with clinically localized PCa^28^ with available transcriptome profiles, the signature of ARE repression was associated with more aggressive tumors (**Fig. 5b**) and was highest in NCCN high and very high risk disease (**Fig. 5b**), and higher Gleason grade, PSA (Extended Data Fig. 10a,b), and stage (Supplementary Table 1 and 2), while ARE activation was associated with the opposite – less aggressive cancers (Fig 5b, Extended Data Fig. 10c-e, Supplementary Table 3 and 4). ARE activation was associated with more luminal features, while the repressive signature was higher in tumors classified as basal subtype (**Fig. 5b**). In a cohort of 855 patients after radical prostatectomy^29^, high expression of the ARE repressive activation signature was associated with worse metastasis-free survival (**Fig. 5c**), while the ARE activated signature higher expression was associated with improved, metastasis-free survival (**Fig. 5d**). Finally, in a cohort of 123 patients with metastatic castration resistant PCa^30,31^, the ARE activation signature was associated with improved overall survival (**Fig. 5e**) and maintenance of luminal character (**Fig. 5f**). Together, these data support that in both clinically localized and advanced, treatment resistant human PCa, genes controlled by AREs are associated with prognosis and luminal versus basal phenotypes.

## Discussion

The transcription factor activity of AR is critical for the development, differentiation, and maintenance of the normal prostate, but “hijacking” of normal AR activity and reprogramming to drive oncogenesis is a fundamental feature of human PCa. The field has made major strides in defining the GAIN of function effects on the AR cistrome and transcriptional program associated with prostate tumorigenesis, including redistribution of AR to enhancers enriched for FOXA1 and HOXB13 motifs^3-5,8,20^. However, the functions of AR that are LOST during this process have been less explored. Here, based on the observation that canonical AREs are depleted in the AR cistromes of human cancers, we designed a novel experimental system to modulate genes specifically controlled by AREs in a genome-wide fashion.

We find that AREs are critical elements for defining distinct gene expression programs controlled by AR, and in particular, mediate differentiation-associated growth suppressive transcription in cells from a prostate lineage. AREs are considered to be a key part of all AR-directed transcription^3,21,32^ – here, however, our data suggests these regulatory elements preferentially control growth suppressive, normal programs, while being relatively dispensable for oncogenic, proliferation associated transcription.

The growth-suppressive nature of AR signaling in normal cells has been a well-known but poorly understood phenomenon at a mechanistic level^33,34^. Others have proposed that a transcriptional repression function of AR mediates this activity^35,36^. Our data showing that direct activation of ARE-containing chromatin regions engages the growth-suppressive effects of AR suggests that selective, context-dependent transactivation is responsible.

The growth-suppressive effects of AR are also highly clinically relevant. We show here that the transcriptional signatures revealed by our ARE modulation strategy are associated with prognosis in human PCa patients, opening up new avenues for biomarker discovery and understanding both prognosis and response to agents targeting AR. Furthermore, activation of AR with supraphysiologic testosterone^37,38^ or cycles of androgen stimulation/deprivation (bipolar androgen therapy) in PCa patients^39-42^ has emerged as promising therapeutic strategies, with clinical trials in progress. This study provides mechanistic insight into these therapeutic approaches and potentially improves the selection of patients for them.

## Methods

### Cell Lines

LNCaP cells (ATCC Item # CRL-1740) were cultured on poly-L-lysine (Sigma-Aldrich; cat. P1274) coated plates in RPMI-1640 containing 10% Fetal Bovine Serum (FBS) and incubated at 37°C and 5% CO2. Cells were passaged twice weekly or when cultures reached 80% confluence. PC3 cells (ATCC CRL-1435) were maintained in DMEM supplemented with 10% FBS, while RWPE-1 cells (ATCC CRL-11609) were maintained in Keratinocyte SFM (1X) medium supplemented with human recombinant epidermal growth factor (rEGF) and bovine pituitary extract. LAPC4 cells were a kind gift from Dr. Robert Reiter (UCLA) and grown in IMDM media containing 10% FBS incubated at 37C. Prostate from the genetically normal mice was harvested at 2-3 months of age and processed and grown as 3D Matrigel culture as previously described^4,13^.

All 2D and 3D cultures were assessed for mycoplasma monthly via the highly sensitive PCR-based kit from ABM (cat. G238). Where applicable, cell line identity was validated yearly through the Human STR profiling cell authentication service provided by ATCC.

### Generation of MACC constructs

The oligos used in the MACC construction are provided in Supplementary Table 5.

#### Cloning AR DBD domain and NLS with ERT2

1. The AR DBD domain was cloned with a mutant estrogen ligand-binding domain (ERT2) to achieve tamoxifen inducibility. The AR DBD domain DNA fragment was amplified using Herculase II Fusion DNA Polymerase (Agilent Technologies; cat. 600677) with 36 cycles and annealing at 63.5°C, then purified with the Qiagen PCR purification kit (Qiagen; cat. 28004).
2. To link the AR DBD domain with ERT2, the pRetroQ-Cre-ERT2 plasmid (Addgene #59701) and AR DBD domain DNA fragment were cut by NheI (N-terminal) and XhoI (C-terminal) and purified with the QIAquick Gel Extraction Kit (Qiagen; cat. 28704). After ligation with T4 DNA Ligase (New England; cat. M0202S), bacterial clone screening and sanger sequencing were performed to verify the AR DBD domain DNA fragment in frame and unmutated. This inducible construct (AR-DBD_ERT2 construct) will be further modified with chromatin structure modulation.

#### Cloning ARE with modulation of chromatin structure

1. To modulate the chromatin structure around the AREs on our inducible construct, we linked repressive or activating chromatin modifying domains (H3K9 methyltransferase KRAB or transcriptional activator VP64) and an epitope tag (3xFlag tag) to the C-terminal of our AR-DBD_ERT2 construct. We amplified 3xFlag-KRAB (KRAB-dCas9 plasmid; Addgene #112195), 3xFlag-VP64 (pSL690 plasmid; Addgene #47753), and 3xFlag (KRAB-dCas9 plasmid; Addgene #112195) fragments using Herculase II Fusion DNA Polymerase (Agilent Technologies; cat. 600677) and primers with 30 cycles and annealing at 63°C, purified them with the Qiagen PCR purification kit (Qiagen; cat. 28004) and verified all products by gel separation and sanger sequencing to ensure they were unmutated.
2. To add the 3xFlag-KRAB, 3xFlag-VP64, and 3xFlag fragments to the C-terminal of our AR-DBD_ERT2 construct, we used NheI-HF (New England Biolabs, cat. R3131L) to cut both the AR-DBD_ERT2 construct and the purified fragments, and Shrimp Alkaline Phosphatase (rSAP) (New England Biolabs, cat. M0371S) after AR-DBD_ERT2 construct NheI fragmentation to dephosphorylate. We then used T4 DNA Ligase (New England; cat. M0202S) to ligate the fragments with a 1:3 insertion to vector ratio at 16C overnight and transformed all reactions into Stbl3 chemically competent cells (Thermo Fisher Scientific; cat. C737303). We performed bacterial clone screening and Sanger sequencing to verify that they were in frame and unmutated.
3. These three inducible constructs (MACC constructs) will be further modified to enable lentiviral production.

#### Cloning the MACC constructs into the lentiviral and doxycycline-inducible backbone

To facilitate lentiviral production, we transferred all MACC constructs into a lentiviral backbone, specifically the pLenti PGK Blast V5-LUC (w528-1) plasmid (Addgene #19166). The process involved using SalI-HF (New England Biolabs, cat. R3138S) and AatII (New England Biolabs, cat. R0117L) to cleave both the MACC constructs and the backbone, followed by ligation, purification, and verification of all products through gel separation and Sanger sequencing to ensure their mutational integrity. To achieve the doxycycline-inducible ability, all the AR DBD domain with modulation of chromatin structure were cloned into a doxycycline-inducible lentiviral backbone, and verification of all products through gel separation and Sanger sequencing was conducted to ensure their mutational integrity.

### Generation of lentivirus

293T cells were cultured in 10 cm tissue culture plates until they reached 70–80% confluency. Transfection was performed using Lipofectamine 3000 Transfection Reagent (Thermo Fisher Scientific; cat. L3000015) with pMD2.G (lentiviral helper plasmid; Addgene #12259), psPAX (lentiviral helper plasmid; Addgene #12260), and the target transfer plasmid. Lentivirus was harvested 48/72 hours after the start of transfection. The lentivirus supernatant from plates transfected with the same plasmid construct was pooled, and cellular debris was removed by filtration using Millipore’s 0.45-µm filter unit. The filtered lentivirus supernatant was aliquoted and stored at −80 °C for later use.

### Generation of stable cell lines

To generate stable cell lines expressing the MACC construct, prostate 2D cells were infected with crude lentivirus at a ratio of 1:500 for 24 hours. 3D organoid lines were generated using a spinoculation protocol at 650g and 32C for 1 hour. Subsequently, the cells and organoids were selected using Blasticidin (InvivoGen; cat. ant-bl-05) for 7 days.

### Xenograft tumor growth

6-8 week old nude male mice were injected subcutaneously with 3 million LNCaP cells stably expressing the neutral, repressive, or activating MACC constructs with 100uL of 1:1 Matrigel (Corning; cat. 354234) and cells (resuspended in 1x PBS) and allowed to grow until they reached a volume of 300mm^3^ on their flank. Mice were fed with normal chow, and then switched to doxycycline-containing chow starting from week 7 until the study’s termination. Tumor volume was measured weekly with electronic calipers and the total volume was calculated using the formula (w^*^l^*^h). Mice were sacrificed if tumor volume surpassed the predetermined upper limits according to the approved IACUC protocol.

### Animal studies approval

All mouse studies were approved by Weill Cornell Medicine (WCM) Institutional Care and Use Committee under protocol 2015–0022.

### Immunoblot

For organoids, protein lysates were prepared after digestion of Matrigel (Corning; cat. 356231) with TrypLE Express Enzyme (Thermo Fisher Scientific; cat. 12605028) and washed in PBS and lysed in RIPA buffer supplemented with protease and phosphatase inhibitors. For cell lines, pelleted cells were washed in PBS and lysed in RIPA buffer supplemented with protease and phosphatase inhibitors. Proteins were quantified by BCA assay and separated on 4%–15% Protein Gels, protein was transferred to a nitro-cellulose membrane using the iBlot semi-dry system from Invitrogen, blocked in 5% milk in TBST, and incubated with primary antibody overnight rocking at 4C. Individual blots were washed 3x in TBST, incubated with species-specific HRP conjugated secondary antibody for 45 min at 24C, washed again 3x with TBST buffer and then imaged using the SuperSignal West Pico PLUS Chemiluminescent Substrate (Thermo Fisher Scientific; cat. 34578) on a ChemiDoc imaging system from BioRad. All antibodies and their concentrations used in this study can be found in Supplementary table 6.

### Immunofluorescence

Cells were plated and grown on poly-L-lysine coated coverslips for 48-72 hours, then fixed with 4% PFA at room temperature for 15 minutes. After washing twice with PBS, cells were permeabilized with 0.1% Triton-X100 in PBS for 10 minutes. Following another wash with PBS, cells were blocked with 10% goat serum and 0.5% BSA in PBS for 30 minutes at room temperature. For organoids, immunofluorescence was performed following the previously described procedures^4,13^. Briefly, the organoids were suspended in Cell Recovery Solution (Corning; cat. 354253) to dissolve the Matrigel while preserving the 3D cellular structure. Next, the organoids were collected and embedded into fibrinogen and thrombin clots. Paraffin sections were prepared at The Translational Research Program of the Department of Pathology and Laboratory Medicine at WCM. Primary antibodies were applied at the specified dilution in blocking solution and incubated overnight at 4°C. The next day, cells were washed twice with PBS and incubated with the appropriate fluorescent secondary antibody (in blocking solution) for 30 minutes in the dark. Coverslips were then washed three times with PBS, mounted with Prolong Gold antifade mount solution with DAPI (Thermo Fisher Scientific; cat. P36931), and visualized using a fluorescent microscope.

### Growth Curves

5000 cells for each cell line were plated in biological triplicate and assessed for changes in confluency over time using Incucyte software (2022B Rev1) for the indicated duration. The mean and standard error of each time point is plotted.

#### Treatments

For androgen stimulation, five thousand cells were plated in a 96-well plate in phenol-red free RPMI-1640 containing 5% charcoal dextran treated serum (HyClone; cat. SH30068.03IR25-40). Cells were allowed to attach overnight and were treated with varying concentrations of DHT (in EtOH). Growth was monitored and calculated using Incucyte software. The average of 4 images per well was plotted in biological triplicate for each cell line and each condition.

For Tamoxifen (Tam) treatment, five thousand cells were plated in a 96-well plate with phenol-red free RPMI-1640 and 10% FBS. Following an overnight attachment period, cells were treated with varying concentrations of Tamoxifen (in EtOH). Confluency was monitored and calculated with Incucyte software. The average of four images per well was plotted in biological triplicate for each cell line and condition. When applicable, Caspase-3/7 Green Reagent for Apoptosis (Sartorius; cat. 4440) was added to the cell culture media, as recommended by the manufacturer, and the fluorescence signal was monitored using an Incucyte live-cell analysis system.

### RNA extraction and library preparation

LNCaP MACC cells and RWPE1 MACC cells were stimulated with either EtOH or Tamoxifen (Sigma-Aldrich; cat. T176) and harvested for RNA-seq in biological replicates. Total RNA was extracted, and DNaseI treated by RNeasy Mini Kit (Qiagen; cat. 74104). Nanodrop quantified RNA was checked by Bioanalyzer RNA 6000 Nano Kit (Agilent Technologies). Samples with RNA integrity number > 10 were used for library preparation (Illumina Stranded mRNA Prep kit) and sequenced on Illumina NovaSeq 6000 at the WCM Genomics Core.

### RNA-seq analysis

Data was processed using (v3.10) of the nf-core collection of workflows^43^. The pipeline was executed with Nextflow^44^ v22.10.4 with the following command:

*nextflow run nf-core/rnaseq -r 3*.*10 --input samplesheet*.*csv --genome GRCh37 -profile singularity*.

Briefly, raw FASTQ files were aligned to the GRCh37 genome and quantified by salmon (v 1.9.0)^45^. Differential gene expression was obtained in R with DESeq2 (1.28.0) package^46^. Gene set enrichment analysis (GSEA)^47^ was run in pre-ranked mode to identify enriched signatures in the Molecular Signature Database (MSigDB). Gene Set Variation Analysis (GSVA)^48^ was used to assess the activity of predefined gene sets in our dataset. GSVA was performed using the R package “GSVA” and the gene sets were obtained from the MSigDB. We calculated the enrichment score for each gene set in each sample using the “gsva” function from the GSVA package with default parameters. The resulting enrichment scores were then used for visualization.

### ChIP and ChIP-sequencing

ChIP was performed as previously described^4^. The LNCaP MACC cells were treated with 2.5uM Tamoxifen for 4 hours or 18 hours. For each replicate, 20 million cells were fixed with 1% formaldehyde for 10 minutes at 24°C, quenched with 0.125 M glycine for 5 minutes, washed 2x with PBS, and stored at -80°C until use. The fixed pellets were lysed in 1% SDS lysis buffer and sonicated for 30 cycles (30s on/off) in a temperature-controlled Bioruptor 3000 to obtain a size range of 250-400 bp. After centrifugation at 15000 RPM for 20 minutes to remove debris, individual samples were incubated with Protein A/antibody-conjugated beads overnight while rocking at 4°C. The samples were washed six times with increasing salt buffers and DNA was eluted at 65°C overnight. All samples were treated with RNase A for 30 minutes at 37°C followed by Proteinase K at 65°C for 1 hour. DNA was purified using phenol chloroform and individual ChIP samples were verified by q-PCR. ChIP sequencing libraries were constructed using the KAPA Hyper Prep Kit (Roche; cat. 08278539001) following the manufacturer’s instructions, using 20 ng DNA per sample. The libraries were assessed for quality, purity, and size using DNA High Sensitivity Bioanalyzer chips (Agilent; cat. 5067-4626), and those passing quality control (with an equal size distribution between 250-400 bp, no adapter contamination peaks, and no degradation peaks) were quantified using the Library Quantification Kit (Roche; cat. 07960298001). Finally, the libraries were pooled to a final concentration of 10nM and sequenced at the WCM Genomics Core on a NovaSeq 6000. The ChIP-seq utilized the ERalpha (ERa) antibody from Santa Cruz Biotechnology (Cat# sc-8002, AB_627558) to generate MACC peaks, along with the H3K27ac antibody from Abcam (Cat# ab4729, AB_2118291).

### ChIP-seq data analysis

Briefly, the quality of the raw reads (FASTQ files) was validated using FastQC software (Version 0.11.7), and single-end reads with a score > 29 were aligned to the hg19 human reference genome using Bowtie2 software (v2.2.9)^49^ with default parameters. The resulting SAM files were converted to BAM format, sorted, PCR duplicates removed, ENCODE blacklist regions eliminated, and final BAM files indexed using Samtools (v1.7)^50^. Replicate BAM files were then combined to generate RPKM-normalized bigwig files for each factor using Deeptools (v3.0)^51^. These bigwig files were used to create heatmaps and binding profiles using Deeptools v3.0 (computematrix, plotProfile, and plotHeatmap functions).

#### Peak calling

Peaks were called using MACS2^52^ with a p value < 10^-8^ or q = 0.05 (replicates combined) using the narrow peak caller and matched input as background. Peak overlap and Venn diagrams were generated using pybedtools and bedtools intersect function^53,54^ and were defined as overlap (more than or equal to) 1 bp. Where indicated, parental AR ChIP-seq data was utilized from GSE117430^4^ and processed and analyzed as above.

A conserved MACC peak set was defined as a peak shared among 2 or more datasets (Neutral, Repressive, or Activating). Overlap between all conditions was determined using bedtools (v2.28.0) intersect function with a minimum overlap of 1bp. Subsets of these peaks were generated using bedtools (v2.28.0) -subtract function.

#### Motif analysis

Motif analysis was performed using Homer software (v4.8.3)^55^ by analyzing a 200 bp window around the center of each peak. Motif density around peaks was calculated using Homer and JASPER^56^ definitions of the conserved motif. To determine motif enrichment between datasets with similar peak numbers, peak sets of the control were used as background (-bg flag in findmotifsgenome.pl function). A p-value less than or equal to 10^-20^ was considered significant for motif enrichment, unless otherwise indicated.

Analyses of AR peaks containing AREs conducted from published ChIP-seq were conducted using peak sets in their published form and the JASPER definition of an ARE. Peaks were assessed for the presence of this motif using HOMER’s annotatepeaks.pl -m function (v3.0) within a 100bp window from the peak center.

The density of specific motifs around a peak set was conducted using the annotatepeaks.pl function in HOMER v3.0 with a window of 2,400bp from the peak center binned every 10 bp. Comparison among ChIP conditions was done by subtracting the determined motif frequency/bp/peak from the other. Signal above 0 was considered enrichment and below 0 depletion. These profiles were plotted in PRISM v9.2.0 and traces were generated using the smooth, differentiate or integrate curve function with 2^nd^ degree curve smoothing. H3K27ac ChIP-seq data from human PCa or normal tissue was downloaded from GSE130408 and analyzed in its published form^20^. Principal component analyses of these samples using conserved MACC ChIP-seq peaks were conducted using DeepTools (v3.0)^51^ MultiBigWigSummary and PlotPCa packages.

### Generation of MACC signatures

Differential expression analyses were performed between tamoxifen and control treatments in each cell (LNCaP MACC cells and RWPE1 MACC cells), and the significantly overexpressed and underexpressed genes were defined as neutral, repressive, and activating signatures. The signature scores were defined as the sum of z-scores from overexpressed genes and underexpressed genes for each signature. Signature score = sum (z-scores from overexpressed genes) – sum (z-scores from underexpressed genes).

### Human data analysis

Human prostate scRNA-seq was downloaded from GSE120716^27^ and analyzed in its published form. De-identified gene expression profiles were obtained prospectively from clinical usage of the Decipher prostate genomic classifier between January 2016 and June 2023, n=169,123 (Veracyte Inc, San Diego, California). Samples were obtained from either prostate biopsy or radical prostatectomy, and ordering criteria for the genomic classifier exclude prior treatment with hormone therapies or radiotherapy. All tumors were prospectively gathered into the Decipher Genomics Resource for Intelligent Discovery (GRID) database (NCT02609269)^28^. A retrospective cohort from individual patient-level data generated in a prior meta-analysis with long-term follow-up (META855; n = 855)^29^ was used to test associations with time to metastasis after radical prostatectomy. Time-to-event end points were shown graphically using the Kaplan– Meier method. Multivariable Cox regressions were used to compare time to failures. The statistical significance of differences in continuous and categorical variables between groups was assessed using Kruskal–Wallis and Pearson X^2^ tests, respectively. Given the exploratory nature, no adjustments for multiple hypothesis testing were performed, all tests were two-sided, and all analyses were performed using R version 4.0.3.

The CRPC cohort consisted of 123 biopsies from male metastatic castration-resistant prostate cancer (mCRPC) patients, derived from two studies^30,31^, with diverse clinical characteristics and treatment histories. Baseline biopsy samples captured initial gene expression at first biopsy after mCRPC diagnosis. RNA-seq aligned with STAR provided gene expression data from gencode v.28, normalized to TPM, and log2 transformed. Z-scores standardized expression values per gene were calculated for ARE activating (VP64) and ARE repressed (KRAB) gene expression scores, summed log2 fold changes for positively and negatively expressed genes. The primary endpoint was overall survival from biopsy to death/last follow-up. Patients were stratified by quartiles of VP64 and KRAB scores. Kaplan-Meier curves visualized survival analysis, and Cox proportional hazards regression evaluated gene expression’s impact on survival as continuous variables per quartile. Basal/luminal gene expression score was calculated as previously described^31^ and was correlated with VP64 and KRAB scores, with Spearman analysis of correlation.

## Availability of materials

All unique biological materials are available from the corresponding author upon reasonable request.

## Data availability

All raw next-generation sequencing, ChIP and RNA–seq data generated in this study have been deposited in the Gene Expression Omnibus (GEO) repository at NCBI under accession code GSE231516. The parental AR ChIP-seq data have been previously published^4^ and are available at the GEO (GSE117430). The H3K27ac ChIP seq data from human PCa or normal tissue have been previously published^20^ and are available at the GEO (GSE130408). The RNA-seq data from TCGA primary prostate cancer patients have been previously published^25^ and are available at the cBioPortal^57^ for Cancer Genomics. The human prostate scRNA-seq data have been previously published^27^ and are available at the GEO (GSE120716).

All other data supporting the findings of this study are in the article, supplementary information or are available from the corresponding author upon reasonable request.

## Acknowledgements

We are indebted to prostate cancer patients and families who contributed to this research. We thank Dr. Dawid Nowak, Dr. Pengbo Zhou, and Dr. Jonathan E. Shoag (WCM) for helpful discussion. We thank the Weill Cornell Medicine (WCM) Genomics Core Facility, the Biospecimen and Pathology Core and the Computational and Biostatistics Core of the WCM SPORE in Prostate Cancer and Memorial Sloan Kettering Cancer Center cBioPortal. This work was funded by: the NCI, US (WCM SPORE in Prostate Cancer, P50CA211024, R37CA215040, and R01CA233650, C.E.B.), Damon Runyon Cancer Research Foundation, MetLife Foundation Family Clinical Investigator Award, US (to C.E.B.), and the Prostate Cancer Foundation, US (M.A.A. and D.L.).

## Author contributions

MAA, XC, and CEB designed the study. MAA, XC and KL designed and executed experiments. DL and AS designed and executed bioinformatic analyses. AH, YL, and ED performed analysis in clinically localized prostate cancer cohorts. MS and FYF performed analysis in CRPC cohorts. FK, ML, BDR provided specimens and performed pathologic analysis. KL, LDD, JTG, and JW provided technical support and performed animal experiments. MAA, XC, DL, and CEB wrote the manuscript. All authors participated in critical evaluation and revision of the manuscript.

