## Supplementary Information for "Canonical AREs are tumor suppressive regulatory elements in the prostate"

**List:**

**Extended data figure 1-10**

**Supplementary figure 1-2**

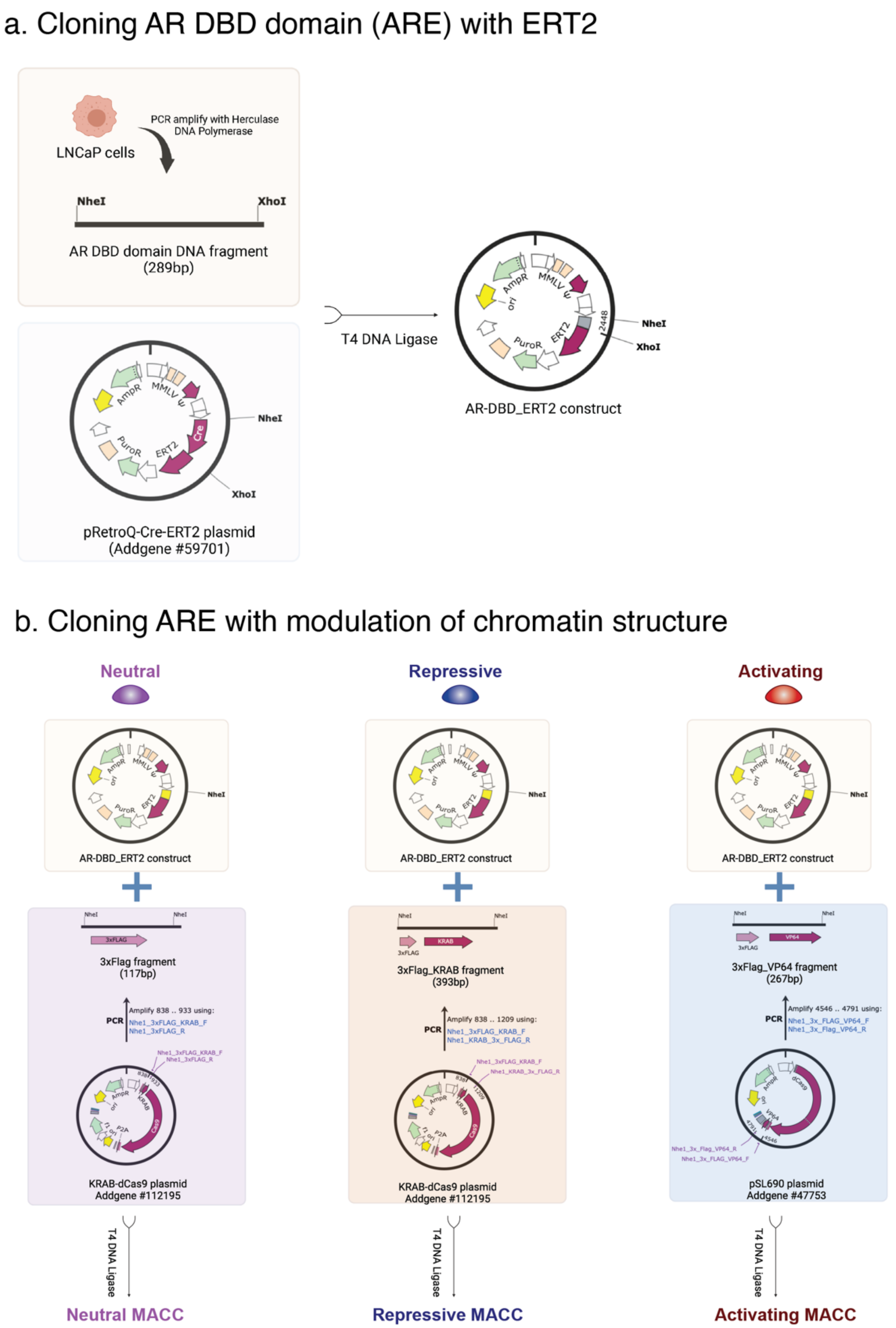


**Extended Data Fig. 1 Cloning strategy of MACC constructs.**
**a,** Cloning AR DBD domain (ARE) with ERT2 to achieve tamoxifen-inducible ability. **b,** Cloning ARE with modulation of chromatin structure.

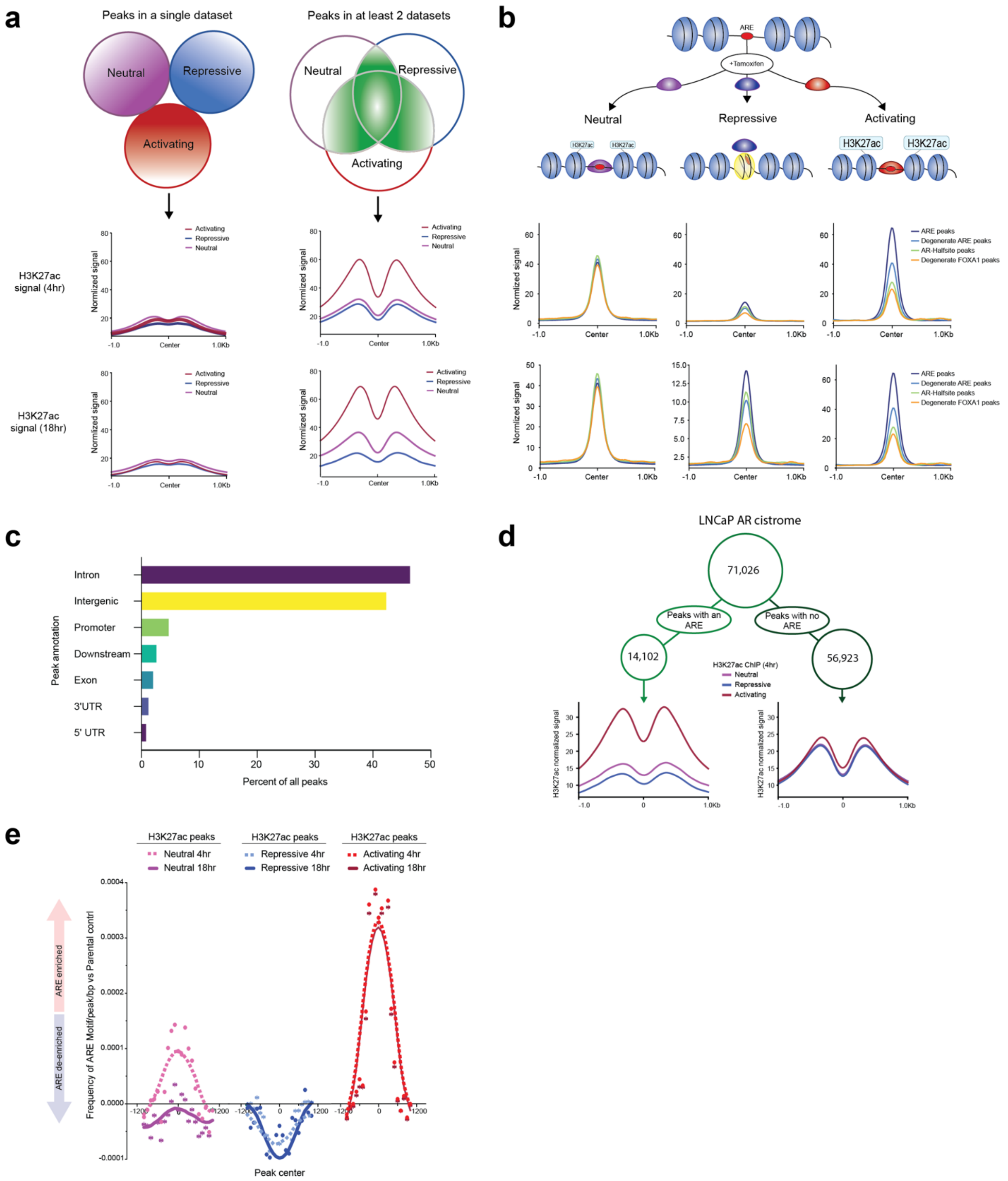


**Extended Data Fig. 2 Generation and validation of consensus MACC ChIP-seq peaks and association with H3K27ac activity.**

**a,** H3K27ac activity of all MACC peaks in different intersections (Neutral, Repressive, and Activating) at 4hr and 18hr. **b,** Peak activity (H3K27ac) distribution of the ARE peaks, degenerate ARE peaks, AR half-site peaks, and degenerate FOXA1 peaks in MACC settings. **c,** Annotation of the MACC peaks by genomic localization as a percentage of all peaks. **d,** Peak activity distribution of LNCaP AR peaks with or without ARE in MACC H3K27ac (4hr) settings. **e,** Changes in ARE motif frequency over parental control of the MACC H3K27ac) peaks (4hr and 18hr.

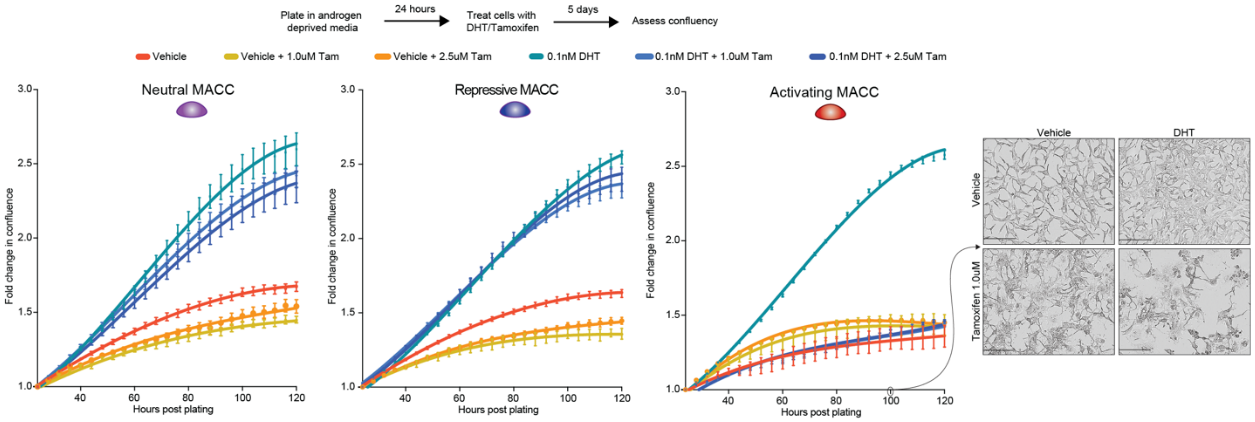


**Extended Data Fig. 3 Growth of LNCaP MACC lines with vehicle or tamoxifen and androgen deprived or DHT stimulated condition.**

The growth of LNCaP MACC lines in 2D culture was assessed after androgen deprivation and DHT stimulation, with the addition of either vehicle or tamoxifen. Cell growth was monitored by live-cell imaging throughout the 5-day treatment period to observe any changes in proliferation rates. Curves represent confluence; representative brightfield images on right.

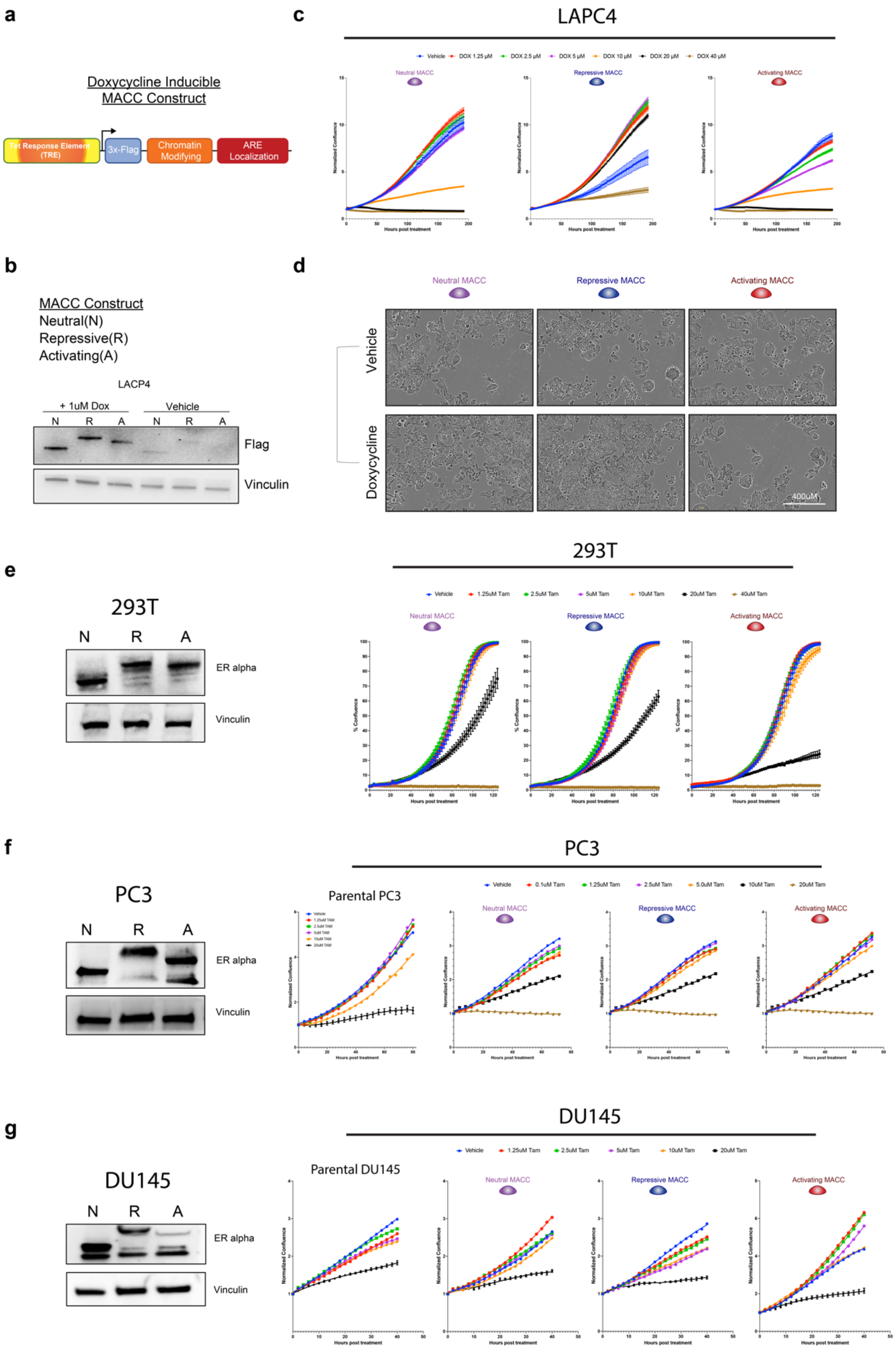


**Extended Data Fig. 4. MACC phenotypic responses are restricted to cell lines of prostate lineage that express AR.**

**a-b,** Schematic of doxycycline inducible MACC constructs, and expression of neutral (3X FLAG only), repressive (KRAB) and activating (VP64) constructs upon doxycycline induction in LAPC4 cells. **c-d,** Growth and brightfield images of LAPC4 MACC lines with vehicle or doxycycline treatment. **e,** Immunoblot and growth of 293T MACC lines with vehicle or increasing doses of tamoxifen treatment. **f-g,** Immunoblot and growth of PC3 and DU145 (two AR-negative prostate cancer cell lines) MACC lines with vehicle or increasing doses of tamoxifen treatment.

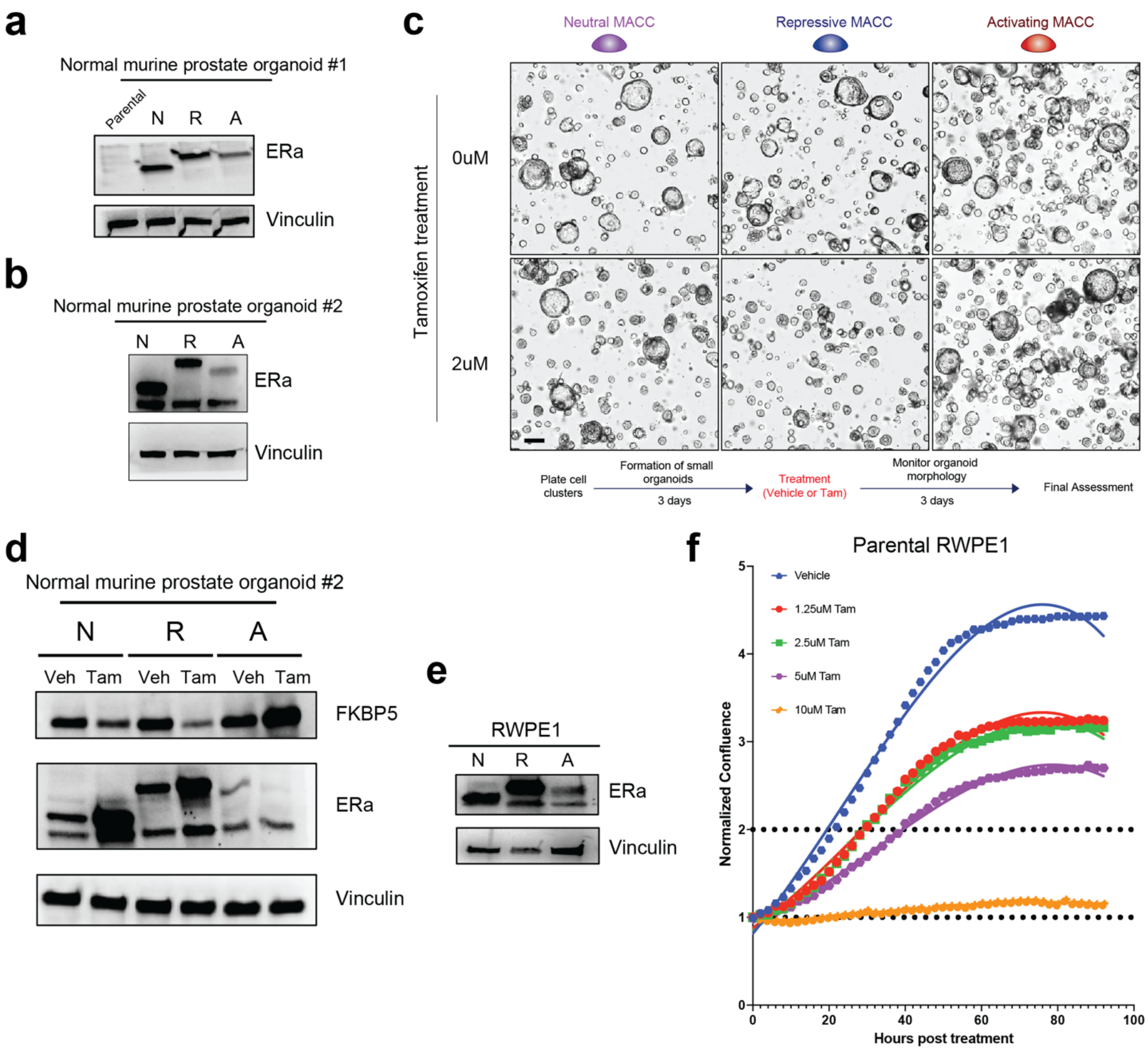


**Extended Data Fig. 5 MACC effects in normal prostate cells.**
**a-b,** Immunoblot of two independent genetically normal mouse prostate organoids with inducible MACC constructs. **c,** Brightfield images and growth of genetically normal mouse prostate organoids with inducible MACC constructs. Scale bar = 40 µm. **d,** Immunoblot of AR target FKBP5 in independent genetically normal mouse prostate organoids with inducible MACC constructs upon vehicle or 2uM tamoxifen treatment. **e,** Immunoblot of RWPE1 prostate lines with inducible MACC constructs. **f,** Growth of parental RWPE1 prostate lines with vehicle or increasing doses of tamoxifen.

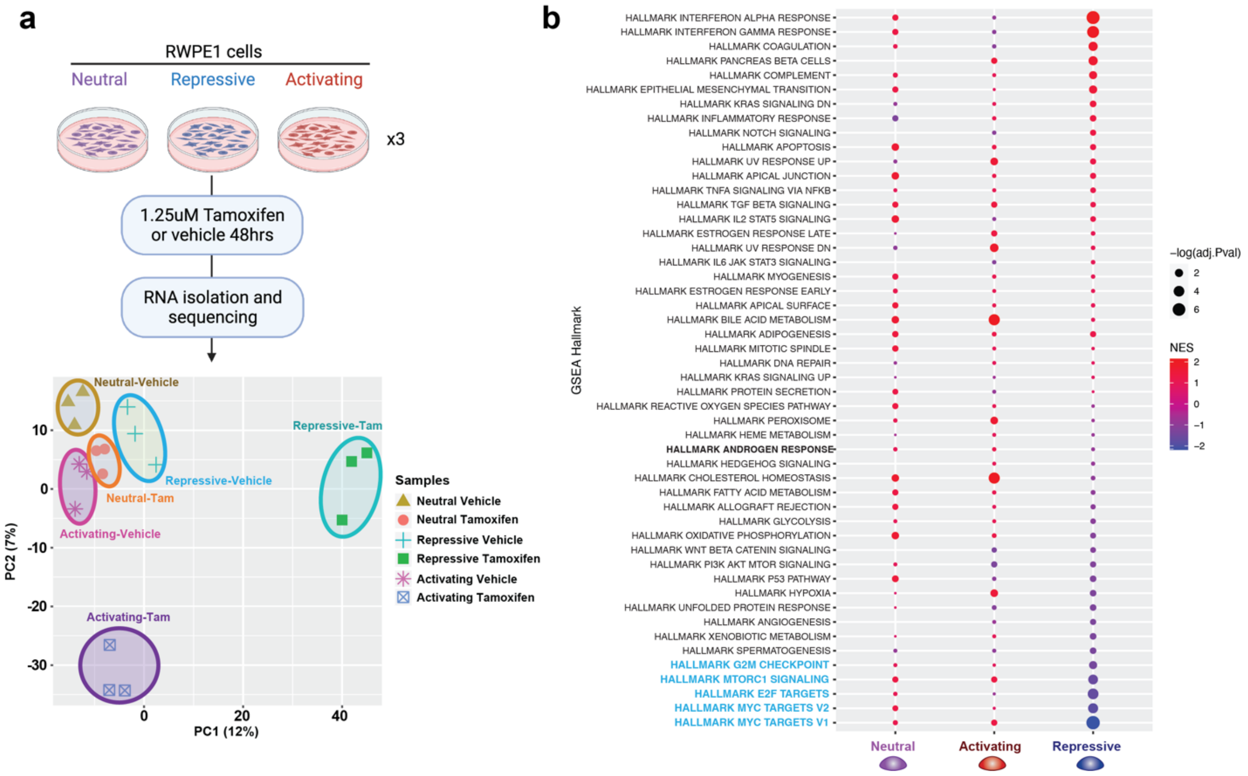


**Extended Data Fig. 6 Transcriptional profiling and pathway analysis of RWPE1 MACC lines.**

**a,** Experimental plan for transcriptional profiling, and PCA of gene expression profiled from vehicle treated or tamoxifen induced RWPE1 MACC lines. **b,** Pathway analysis using GSEA Hallmarks of induced vs. vehicle RWPE1 MACC lines. **a**, Created with BioRender.com.

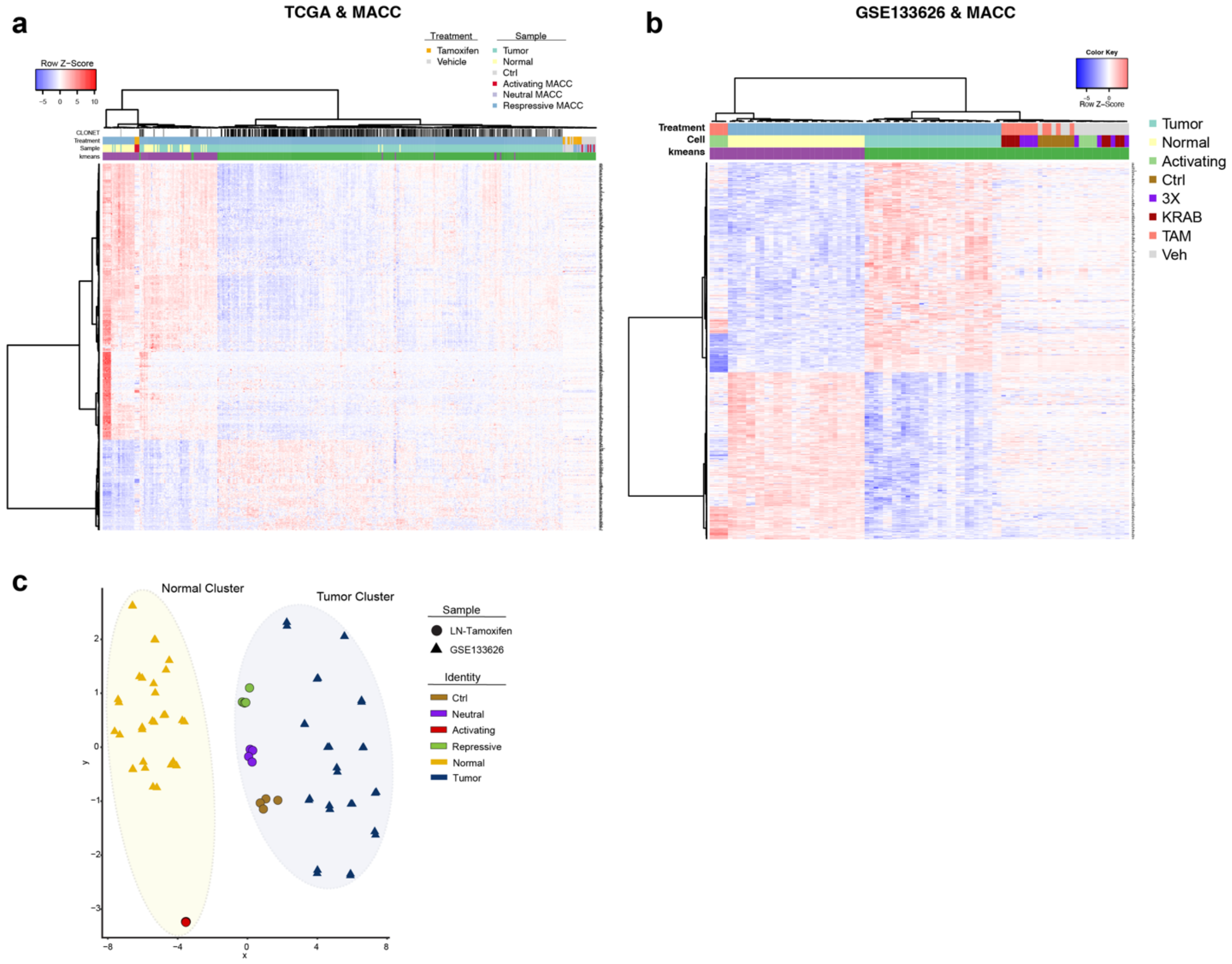


**Extended Data Fig. 7 Unsupervised clustering of the transcriptomes of the human PCa cohorts with MACC.**

**a,** Heatmap of unsupervised clustering of the transcriptomes of TCGA cohort, along with tamoxifen induced LNCaP MACC lines. **b,** Heatmap of unsupervised clustering of the transcriptomes of GSE133626 cohort, along with tamoxifen induced LNCaP MACC lines. **c,** Unsupervised clustering of the transcriptomes of human prostate cancer and normal samples from matched patient (GSE133626), along with tamoxifen induced LNCaP MACC lines. Induced activating MACC lines cluster with normal samples.

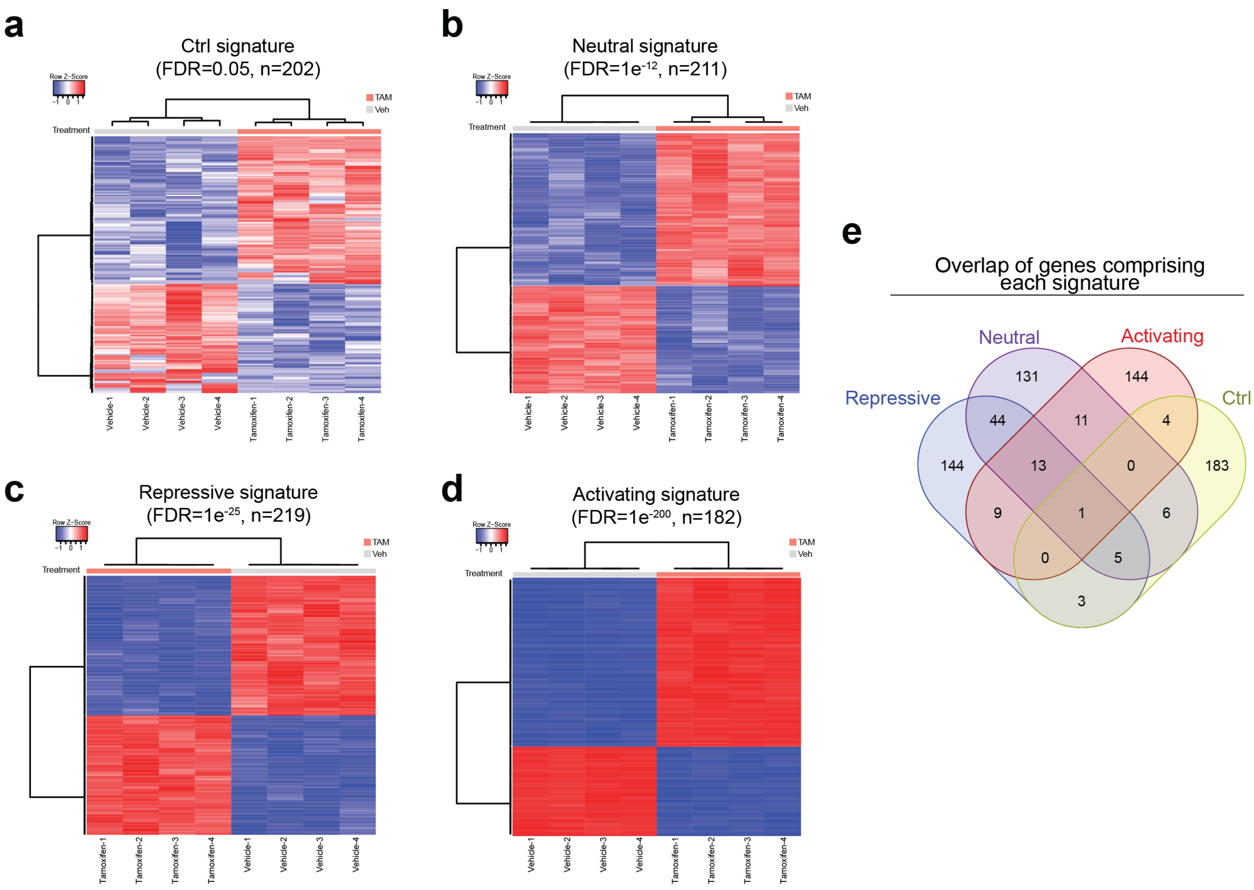


**Extended Data Fig. 8 Generation of MACC signatures using LNCaP models.**
**a-d,** Heatmap of MACC gene signatures from LNCaP ctrl (**a**) cells stimulated with vehicle or tamoxifen, LNCaP Neutral (**b**) cells, LNCaP Repressive (**c**) cells and LNCaP Activating (**d**) cells. **e,** Venn plot of overlapping all MACC gene signatures.

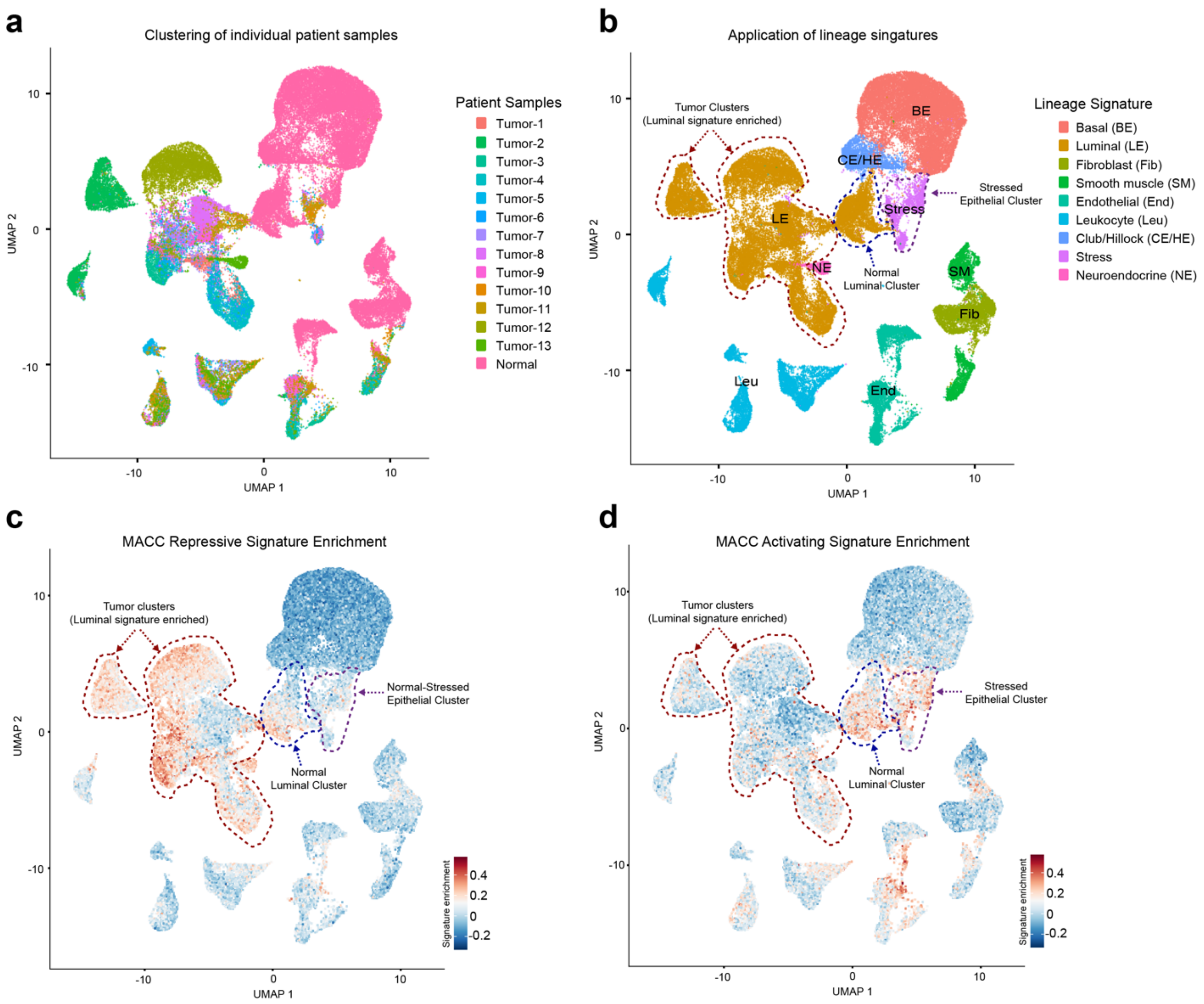


**Extended Data Fig. 9 Enrichment of MACC signatures in scRNA-seq data from tumor and normal prostate samples**

**a,** Uniform Manifold Approximation and Projection (UMAP) view of single cells, color-coded by patient samples. **b,** UMAP view of single cells, color-coded by assigned cell type. **c-d,** UMAP view of single cells by expression level of MACC Repressive (c) and Activating (d) signatures.

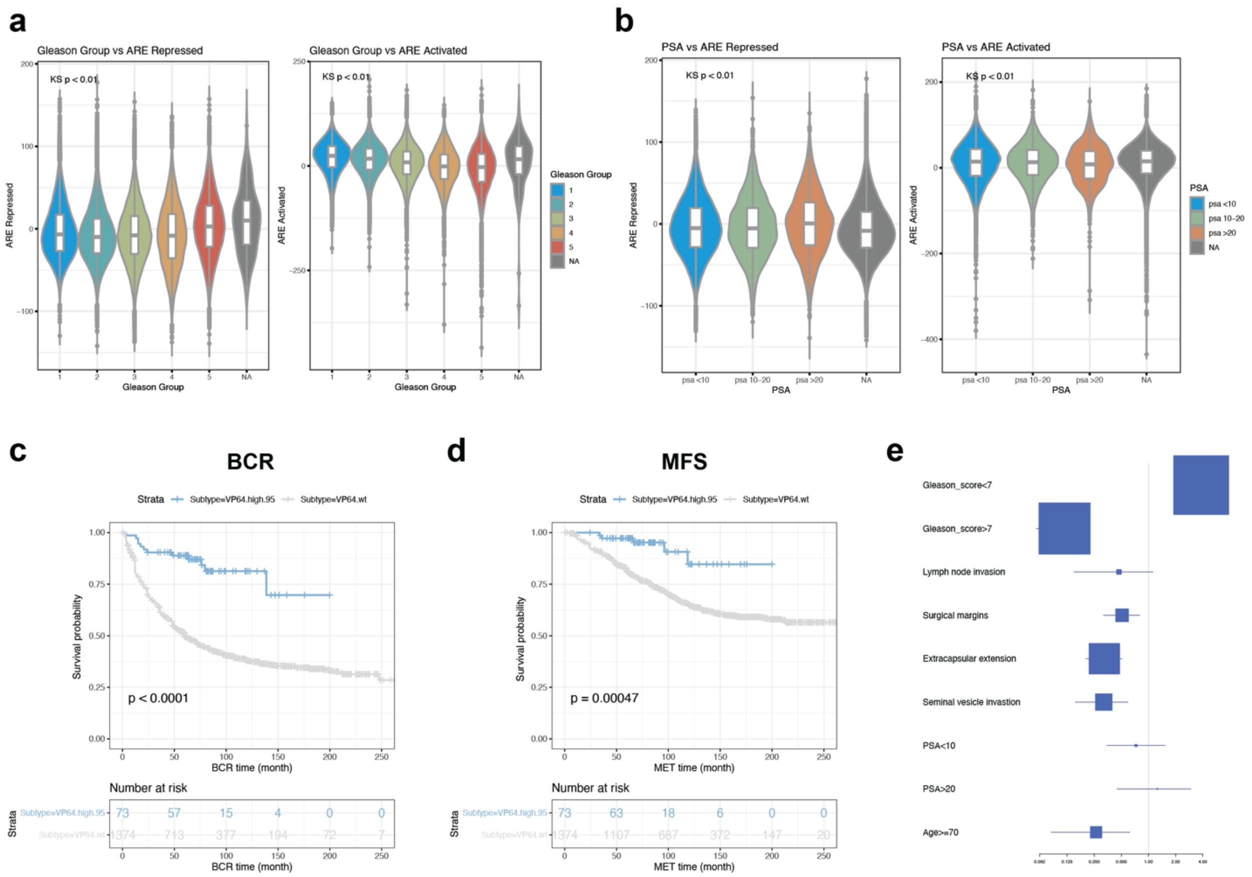


**Extended Data Fig. 10 MACC transcriptional programs are clinically relevant.** **a-b,** Association of ARE activated and ARE repressed signatures from LNCaP MACC lines with Gleason group (**a**) and PSA level (**b**) in clinically localized prostate cancer, n=169,123. KS = Kruskal-Wallis. **c-d,** Kaplan Meier analysis of the biochemical recurrence (BCR, **a**) and metastasis-free survival (MFS, **b**) in a pooled retrospective cohort (n=1677), stratified by ARE activated (VP64) signature expression level. Time = months. **e,** Clinical and pathological associations of the ARE activated signatures from LNCaP MACC lines in the Decipher cohort via univariate analyses.

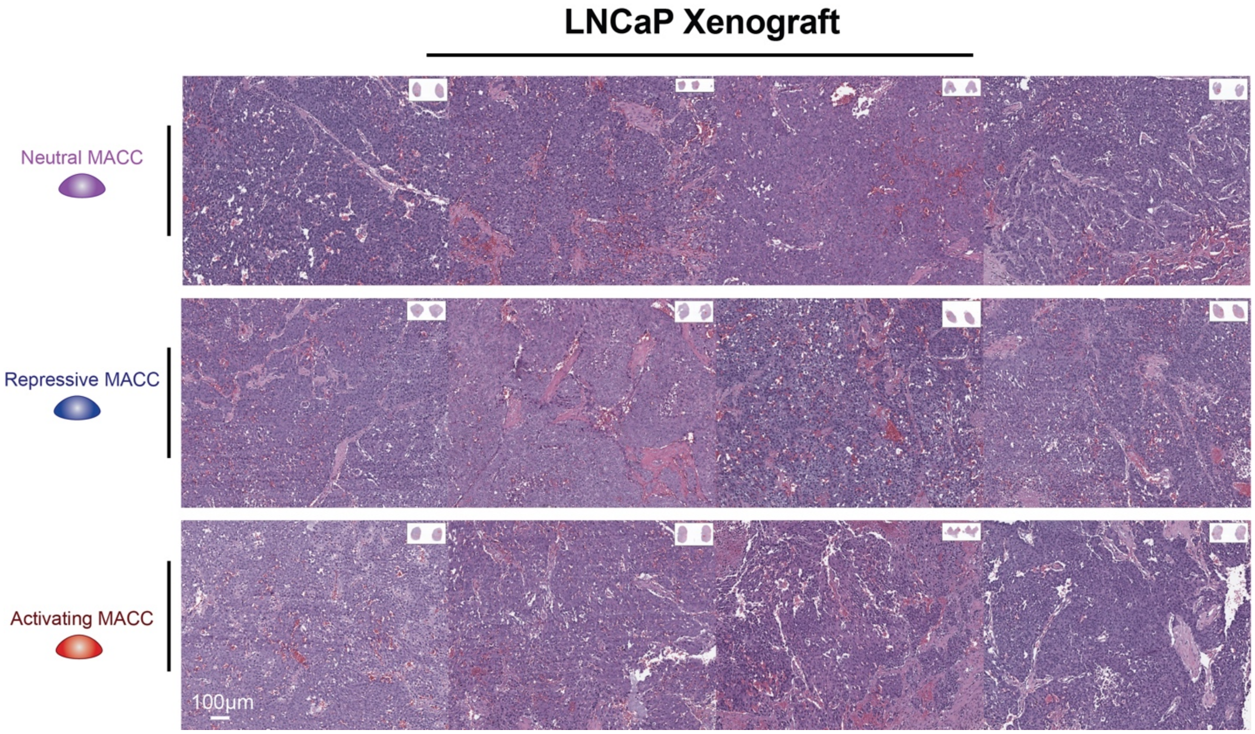


**Supplementary Fig.1 H&E staining of LNCaP xenograft (scale bar 100 um).**

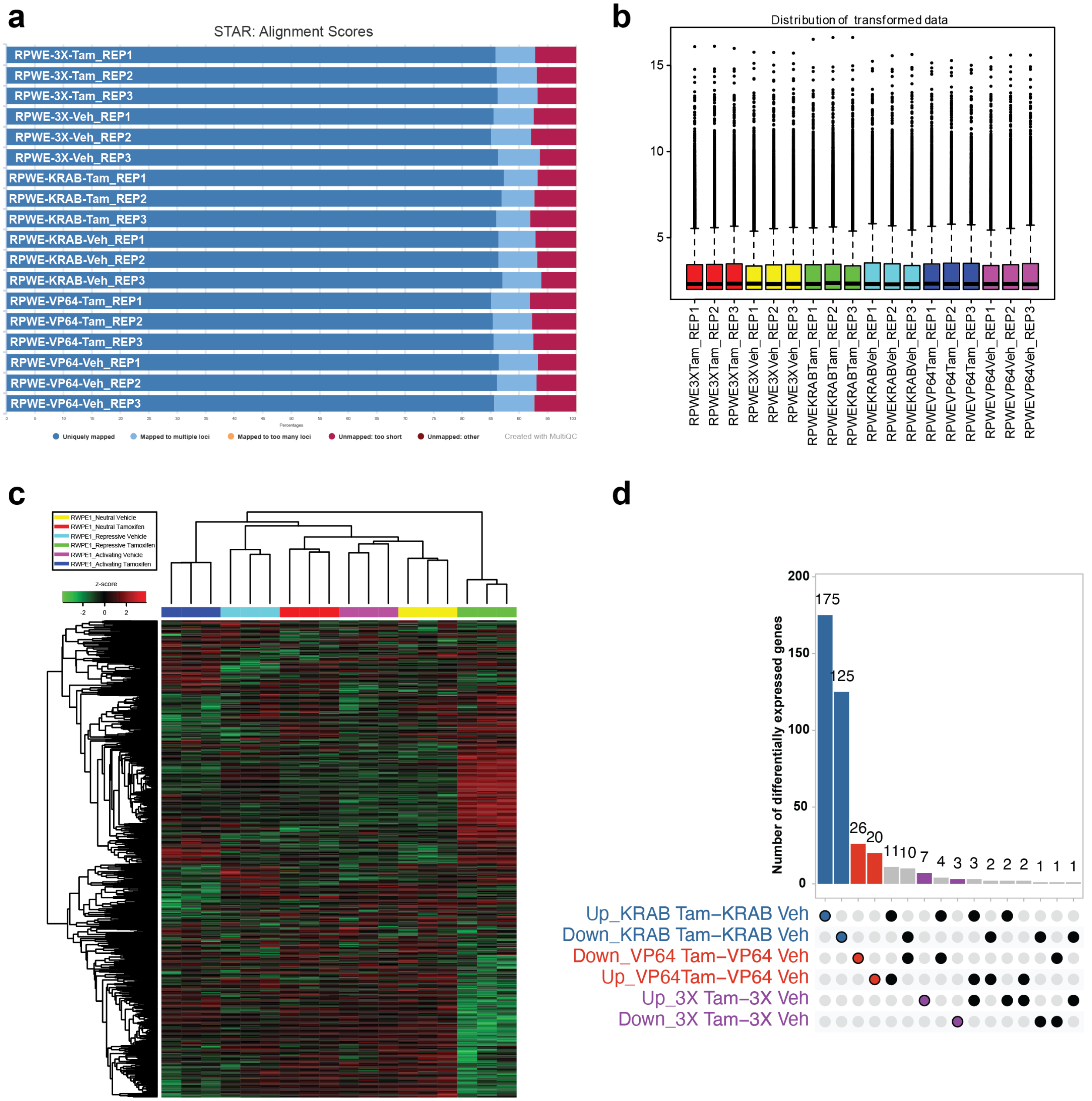


**Supplementary Fig.2 Quality control of RWPE1 MACC RNA-seq.**
**a,** Alignment score of mapping RNA-seq reads to the reference genome. **b,** The distribution of read counts. **c,** The heatmap of the gene expression profiles of different sample groups within the RWPE1 MACC RNA-seq dataset. **d,** Upset plot of differentially expressed genes among the various groups in the RWPE1 MACC RNA-seq analysis.

**Supplementary Table 1 ARE repression score with clinical characteristics in localized prostate cancer patient cohort.**

**Supplementary Table 2 Univariable and multivariable Cox regression analyses of ARE repression score.**

**Supplementary Table 3 ARE activation score with clinical characteristics in localized prostate cancer patient cohort.**

**Supplementary Table 4 Univariable and multivariable Cox regression analyses of ARE activation score.**

**Supplementary Table 5 Oligos or DNA sequences used in this study.**

**Supplementary Table 6 Antibodies used in this study.**
